# Actin filaments accumulate in the nucleus during interphase and remain in the vicinity of condensing chromosomes during prophase to metaphase in zebrafish early embryos

**DOI:** 10.1101/2022.11.06.515321

**Authors:** Haruka Oda, Yuko Sato, Shigehiro A. Kawashima, Yusuke Fujiwara, Máté Pálfy, Edlyn Wu, Nadine L. Vastenhouw, Motomu Kanai, Hiroshi Kimura

## Abstract

In the cytoplasm, filamentous actin (F-actin) plays a critical role in cell regulation, including cell migration, stress fiber formation, and cytokinesis. Recent studies have shown that actin filaments that form in the nucleus are associated with diverse functions. Here, using live imaging of an F-actin-specific probe, superfolder GFP-tagged utrophin (UtrCH-sfGFP), we demonstrated the dynamics of nuclear actin in zebrafish (*Danio rerio*) embryos. In early zebrafish embryos up to around the high stage, UtrCH-sfGFP increasingly accumulated in nuclei during the interphase and reached a peak during the prophase. After nuclear envelope breakdown (NEBD), patches of UtrCH-sfGFP remained in the vicinity of condensing chromosomes during the prometaphase to metaphase. When zygotic transcription was inhibited by injecting α-amanitin, the nuclear accumulation of UtrCH-sfGFP was still observed at the sphere and dome stages, suggesting that zygotic transcription may induce a decrease in nuclear F-actin. The accumulation of F-actin in nuclei may contribute to proper mitotic progression of large cells with rapid cell cycles in zebrafish early embryos, by assisting in NEBD, chromosome congression, and/or spindle assembly.

**Summary statement:** Filamentous actin accumulates in the nucleus of zebrafish early embryos and forms patches associating with condensing chromosomes during prophase.

## Introduction

Actin is a highly conserved protein throughout eukaryotes. In the cytoplasm, actin serves a role in cell migration, stress fiber formation, and cell division by forming a contractile ring during cytokinesis. Actin is also present in nuclei, where it plays a crucial role in transcriptional regulation (Vartiainen *et al*., 2007; Baarlink *et al*., 2013; Tian *et al*., 2016; Sokolova *et al*., 2018) and in DNA damage repair processes (Chiolo *et al*., 2011; Belin *et al*., 2015; Caridi *et al*., 2019). Actin nuclear localization is mediated through the interplay of nuclear import regulated by importin 9 with cofilin (Dopie *et al*., 2012) and nuclear export regulated by exportin 6 (Exp6), which transports profilin-actin complexes (Stüven *et al*., 2003). Whereas filamentous actin (F-actin) is rarely observed in somatic cell nuclei, it is presumed to be involved in regulating the function and structure of undifferentiated cell nuclei (Misu *et al*., 2017).

In mouse early embryonic nuclei, F-actin facilitates nuclear expansion during the G1 phase (Baarlink *et al*., 2017) and maintains the integrity of the totipotent state of the cell mass (Okuno *et al*., 2020). Nuclear actin polymerization via Wave-1 is required for reprogramming somatic nuclei that are transplanted into African clawed frog (*Xenopus laevis*) oocytes (Miyamoto *et al*., 2011). Actin has a distinct role in the large nucleus of oocytes, which is also called the germinal vesicle (GV). *Xenopus laevis* GV contains a high concentration of actin to maintain the large nuclear structure (Clark and Merriam, 1977; Clark and Rosenbaum, 1979; Bohnsack *et al*., 2006). In mammalian oocytes, F-actin bundle in GVs stabilizes the chromatin mobility and secures the accurate chromosome alignment and segregation (Mogessie and Schuh, 2017; Scheffler *et al*., 2022; Dunkley *et al*., 2022). In starfish oocytes, actin mesh forms in the nuclear area after nuclear envelope break down (NEBD; Lénárt *et al*., 2005), and the contraction of the actin meshwork transports chromosomes until they are captured by microtubules (Mori *et al*., 2011). In addition, F-actin polymerizes underneath the nuclear envelope and accelerates fragmentation of the nuclear envelope during the first stage of NEBD (Mori *et al*., 2014; Wesolowska *et al*., 2020).

During *Xenopus* embryonic development, F-actin accumulates in nuclei up to the blastula stage, when the major wave of zygotic transcription begins, before diminishing at the gastrula stage (Oda *et al*., 2017). A reconstituted system utilizing *Xenopus* egg extracts has been used for analyzing the assembly of early embryonic nuclei from de- membraned sperm *in vitro* (Murray, 1991). In reconstituted nuclei using this system, nuclear actin prevents chromatin aggregation during interphase and facilitates chromosome alignment during prometaphase (Oda *et al*., 2017). However, the dynamics of nuclear actin during *Xenopus* embryo development have not been elucidated, because live-cell imaging is hampered by the melanin granules that are present in the cortex of embryos.

In contrast to the African clawed frog (*Xenopus laevis*), zebrafish (*Danio rerio*) embryos that share a similar developmental process with frog are transparent and thus suitable for live-cell imaging. During the early embryo stages of zebrafish, cells have relatively large nuclei and undergo a series of short and synchronized cell cycles without transcription and gap phases (Satoh, 1977; Kimmel *et al*., 1995; Joseph *et al*., 2017), which also occurs in *Xenopus* embryos (Jevtić and Levy, 2015; Newport and Kirschner, 1982; Lucas *et al*., 2000; Mahbubani *et al*., 1992).

In zebrafish embryos, transcription from the zygotic gene is observed as early as at the 64-cell stage, and becomes further activated at around the 512-cell to 1k-cell stages (Mathavan *et al*., 2005; Heyn *et al*., 2014; White *et al*., 2017; Pálfy *et al*., 2017; Vastenhouw *et al*., 2019). After the 1k-cell stage, the cell cycle elongates and loses synchronicity, which occurs with the onset of differentiation (Kane *et al*., 1992).

Fluorescently labeled phalloidin has been the standard reagent to stain F-actin in fixed cells (Dancker *et al*., 1975; Wehland *et al*., 1977). Live-cell actin-specific probes have also been developed, including fluorescent protein-tagged LifeAct and UtrCH, as well as actin-specific chromobody (Actin-chromobody®, Rocchetti *et al*., 2014). UtrCH, which is the Calponin Homology domain of human utrophin (Burkel *et al*., 2007), is an actin-binding peptide that is more selective to F-actin than LifeAct, which is derived from the yeast actin-binding protein, Abp140 (Riedl *et al*., 2008). In this report, we mainly used superfolder GFP-tagged UtrCH (UtrCH-sfGFP) to track nuclear F-actin dynamics in zebrafish early embryos. We observed that F-actin accumulates in nuclei during interphase and the actin patches remained in the vicinity of condensing chromosomes during the prophase to metaphase.

## Results

### Visualizing nuclear F-actin in zebrafish early embryos

To investigate whether F-actin accumulates in nuclei in zebrafish early embryos, we first aimed to detect F-actin by using fixed embryos stained with fluorescently labeled phalloidin. DNA was counterstained with Hoechst dye. Phalloidin signals were enriched in nuclei in the 32-, 128-, and 1k-cell stages, particularly in Hoechst-poor regions (Fig. 1A-C). In the sphere stage embryos, the total nuclear phallodin signals were decreased whereas filamentous structures were observed (Fig. 1B and 1C). This observation suggests that F-actin is abundantly present in nucleoplasm during the early stages of zebrafish development, as was observed in *Xenopus laevis* embryos (Oda *et al*., 2017). Although the filamentous structures observed at the later stages are of interest, we here focused on the dynamics and function of nuclear F-actin that are abundantly accumulated at the earlier stages.

**Figure 1.**
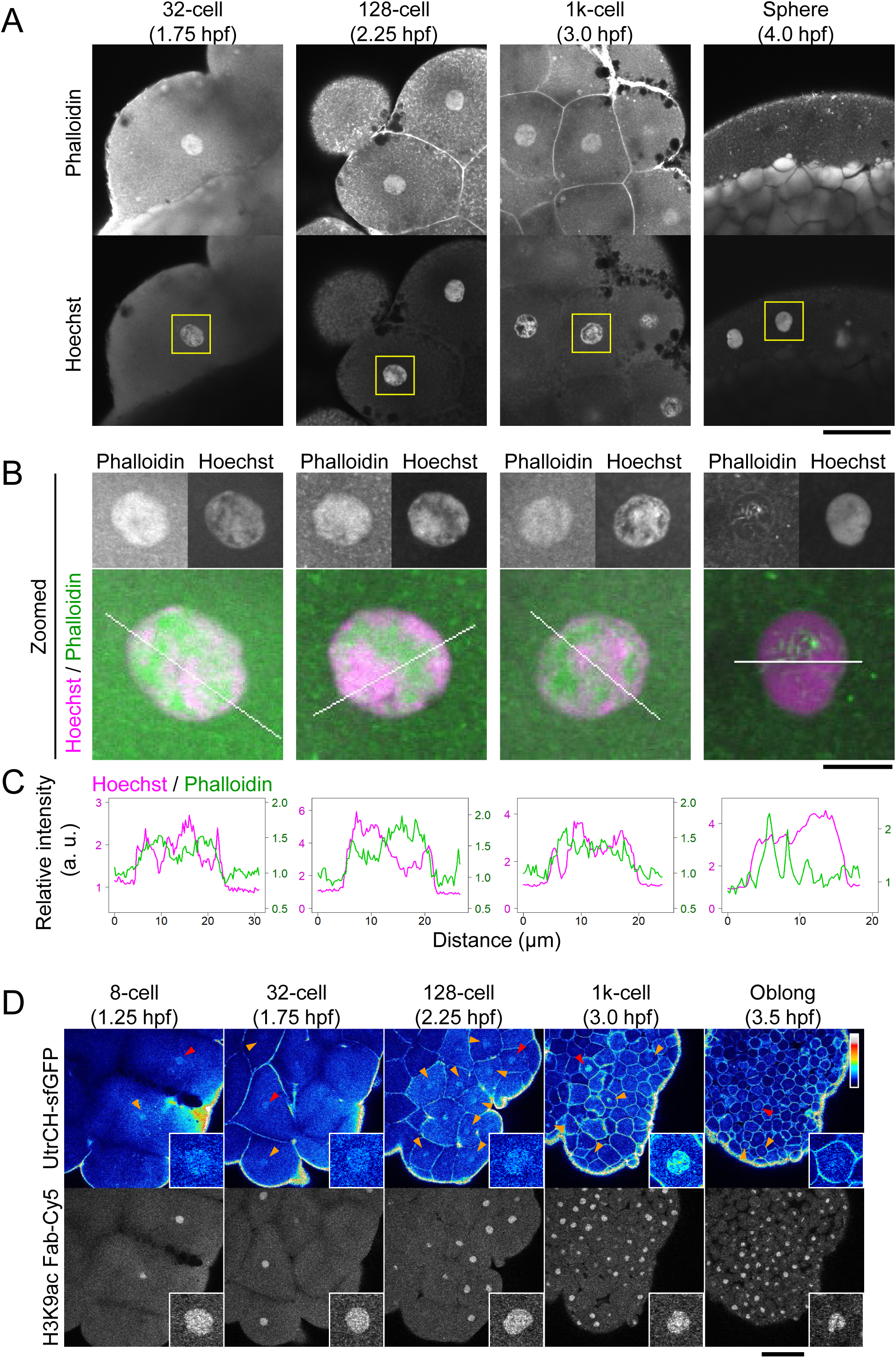
Nuclear F-actin in zebrafish early embryos detected by specific probes. (**A**-**C**) F-actin in fixed zebrafish embryos. Zebrafish embryos were fixed at the 32-, 128-, 1k-cell, and sphere stage, and stained with Acti-stain™ 555 phalloidin and Hoechst. Low (**A**) and high (**B**) power views of single confocal sections are shown. (**C**) Relative intensity profile plots of lines indicated in (**B**). The intensity is normalized using the average cytoplasmic intensity. Phalloidin signals were clearly observed in nuclei of the 32-, 128-, and 1k-cell stages. (**D**) F-actin in living zebrafish embryos. Zebrafish embryos were injected with UtrCH-sfGFP as an F-actin probe and Cy5-labeled Fab for H3K9ac as a chromatin marker. Every 90 s, 25 z-stack fluorescence images with 4 μm intervals were acquired using a confocal microscope. Single z-sections are displayed. Accumulation of UtrCH-sfGFP in nuclei is clearly observed in the 8- to the 1k-cell stage embryos (arrowheads). Insets show the zoomed images of nuclei indicated by red arrowheads. See also Movie 1. Scale bars: 50 μm (**A**), 10 μm (**B**), and 100 μm (**D**).

To visualize nuclear F-actin dynamics in living zebrafish embryos, purified UtrCH-sfGFP (Fig. S1A) was injected into the yolk of 1-cell stage embryos. The antigen-binding fragment (Fab) specific for acetylation of histone H3 lysine 9 (H3K9ac), which distributes throughout the chromatin from the early stage of development, labeled with Cy5 was co-injected as a chromatin marker (Sato *et al*., 2019). The injected embryos were set onto a confocal microscope at the 4-cell stage, and three-dimensional time-lapse images were acquired every 90 s from the 8-cell stage to the oblong stage. UtrCH-sfGFP signal is enriched in the interphase nuclei of embryos from the 8-cell stage to the 1k-cell stage (Fig. 1D, arrowheads and insets; Fig. 2A; Movie 1). At the oblong stage (3.5 hours post-fertilization; hpf), UtrCH signals in nuclei became weaker compared to those in earlier stages (Figs. 1D). These live-cell imaging data are consistent with the observations in fixed embryos.

**Figure 2.**
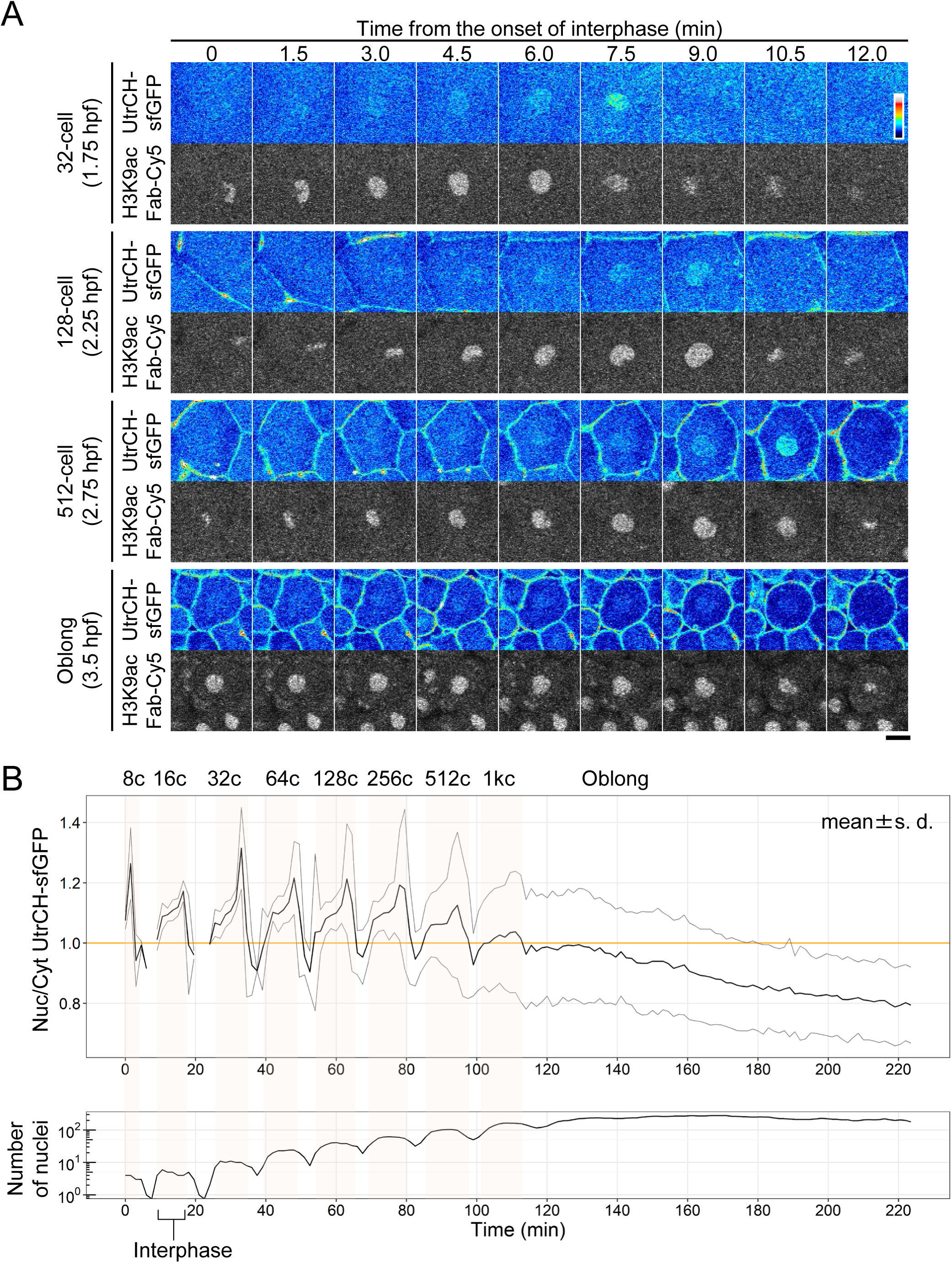
Nuclear F-actin levels increased in the late interphase. Zebrafish embryos were injected with UtrCH-sfGFP and H3K9ac Fab-Cy5, and images were acquired using the same procedure as described in Figure 1D. (**A**) Single confocal sections of a nucleus in different cell stages are depicted with the time (min) after the onset of the interphase. Scale bar: 20 μm. (**B**) The nucleus to cytoplasmic (N/C) intensity ratios of UtrCH-sfGFP were measured. The mean values of UtrCH-sfGFP N/C ratios with the standard deviations (s. d.) and the number of nuclei, as identified by H3K9ac Fab- Cy5, are plotted. The duration of interphase is shaded in orange from the 8- to 1k-cell stage. N/C ratios of UtrCH-sfGFP are above 1.0 (orange line) up to the 1k-cell stage.

Time-laps imaging with 1.5-min intervals (Fig. 2A and Fig. S2) and measurements of the nucleus to the cytoplasm intensity ratio (N/C ratio) of UtrCH-sfGFP (Fig. 2B) revealed that UtrCH-sfGFP signal began to accumulate in the nucleus at the end of telophase, and became more concentrated during interphase, peaking just before mitosis, and then disappeared from mitotic chromosome areas during 16-cell to 1k-cell stages. After the 1k-cell stage, the mean value of UtrCH-sfGFP N/C ratio reduced to less than 1.0 (Fig. 2B).

Because UtrCH could stabilize actin filaments and influence actin dynamics (Spracklen *et al*., 2014), we used another actin-binding peptide, LifeAct, which binds to globular actin (G-actin) and F-actin (Riedl *et al*., 2008), as the sfGFP-tagged version (LifeAct-sfGFP) to validate the results obtained with UtrCH (Fig. 2). Purified LifeAct-sfGFP (Fig. S1B) was co-injected with Cy3-labeled anti-phosphorylated histone H3 serine 28 (H3S28ph) Fab to highlight chromosomes from the prometaphase to anaphase (Hayashi-Takanaka *et al*., 2014) and Cy5-labeled anti-H3K9ac Fab to label nuclei (Fig. S1C). LifeAct-sfGFP was concentrated in nuclei, showing >1.0 N/C ratio, during early embryonic stages (Fig. 3). After the 1k-cell stage, the LifeAct-sfGFP N/C ratio reduced to <1.0 (Fig. 3B), as was observed in UtrCH-sfGFP (Fig. 2B). This lowered N/C ratio on average could be attributed to the less nuclear accumulation and/or the asynchronized cell cycle, as cells in the mitotic phase showed a low N/C ratio (Figs. 2 and 3).

**Figure 3.**
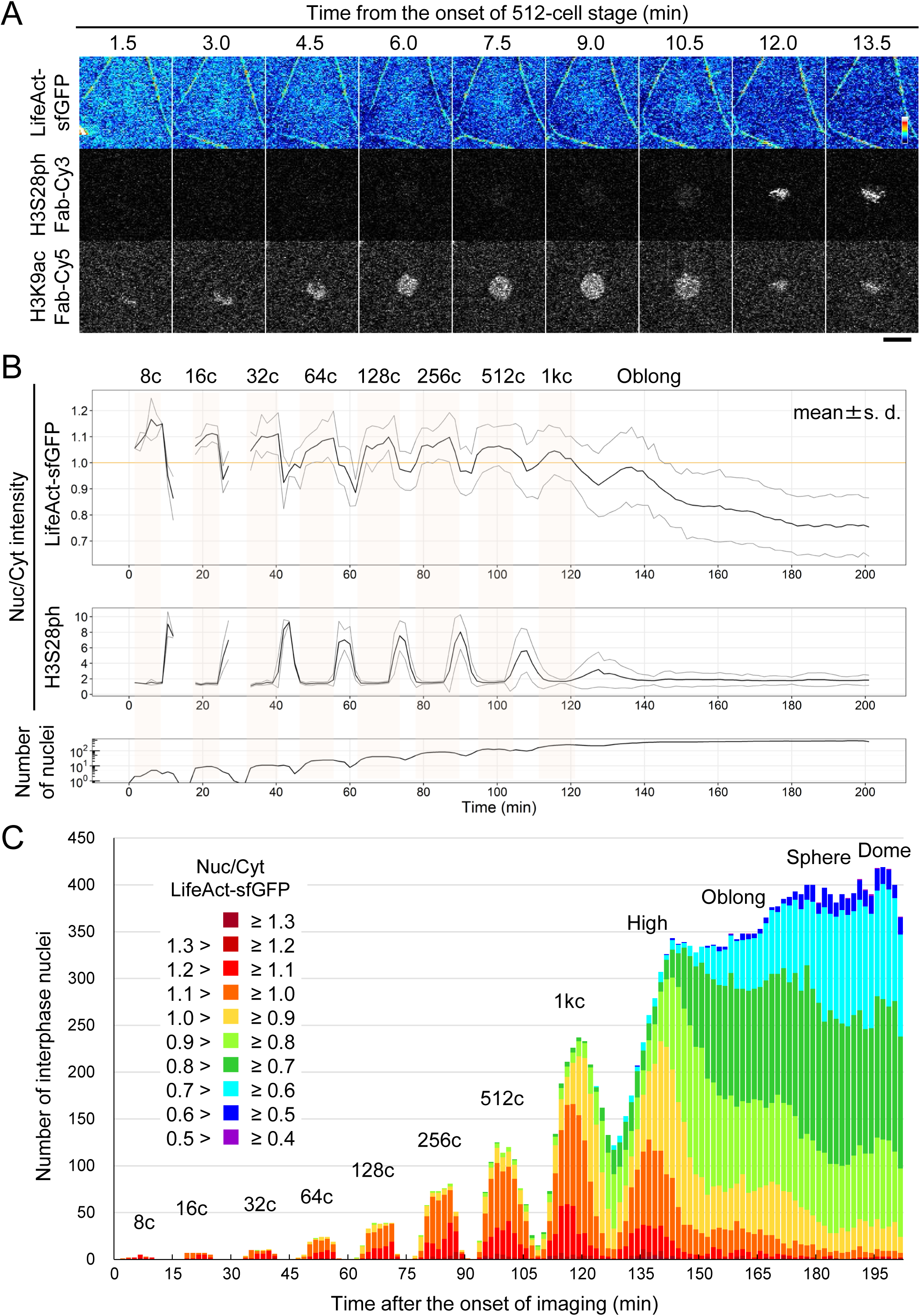
LifeAct-sfGFP accumulates in nuclei in early stage embryos. Zebrafish embryos were injected with LifeAct-sfGFP as an actin probe, H3S28ph Fab- Cy3 as a mitotic chromosome marker, and H3K9ac Fab-Cy5 as a chromatin marker. Every 90 s, 25 z-stack fluorescence images with 4 μm intervals were acquired using a confocal microscope. (**A**) Single confocal sections of a nucleus at the 512-cell stage (2.75 hpf) are shown. LifeAct-sfGFP accumulated in the nucleus up to 12 min and disappeared at 13.5 min, when H3S28ph was observed on condensed chromosomes. Scale bar: 20 μm. (**B**) Mean values of N/C ratios are plotted with standard deviations (s. d.). The duration of interphase is shaded in orange from the 8- to 1k-cell stage. N/C ratios of LifeAct-sfGFP are above 1.0 (orange line) up to the 1k-cell stage. (**C**) Interphase nuclei were selected based on a low H3S28ph Fab signal (N/C ratio <2.2). The numbers of nuclei with different ranges of LifeAct-sfGFP N/C ratio in every time point are plotted as cumulative bars. After the 1k-cell stage, the number of nuclei with LifeAct-sfGFP N/C ratio <1.0 increased.

We then analyzed N/C ratios in individual interphase nuclei by eliminating H3S28ph- positive mitotic cells (Fig. 3C). The cumulative bar chart showed that >90% of interphase nuclei exhibited a LifeAct-sfGFP N/C ratio >1.0 up to the 256-cell stage. At the 512- and 1k-cell stages, the fraction of nuclei with an N/C ratio of >1.0 decreased to ∼80% and ∼65%, respectively (Fig. 3C). After the 1k-cell stage, the fraction of nuclei with an N/C ratio of <1 increased substantially (Fig. 3C, 150 min), reaching ∼90% at the sphere (180 min) and dome (195 min) stages. Taken together with the phalloidin-staining and UtrCH- sfGFP data, these results suggest that actin is gradually concentrated and polymerized in nuclei during the interphase in the early stages of zebrafish embryos, as observed in *Xenopus* (Oda *et al*., 2017), and the amount of nuclear actin decreases after the 1k-cell stage.

### F-actin accumulates in the nucleus during interphase to prophase and remains along with condensing chromosomes during early prometaphase

Since nuclear UtrCH-sfGFP reached a peak just before mitosis (Fig. 2A, 7.5, 9.0, and 10.5 min; Fig. S2A, 7.5, 9.0, and 10.5 min, 3A, 10.5 min), we investigated F-actin dynamics during the late interphase to mitosis in detail. For this purpose, tetramethylrhodamine-labeled 155-kDa dextran (TMR-dextran) was used for monitoring the timing of NEBD; TMR-dextran that was injected into the cytoplasm cannot enter to the nucleus due to its big size until the nuclear membrane breaks down at the onset of prometaphase. Thus, NEBD was identified by the onset of TMR-dextran leakage into the nucleus. In addition, we used JF646-labeled latency-associated nuclear antigen peptide (JF646-LANA) to track chromatin throughout the cell cycle. The LANA peptide binds to the acidic patch in the nucleosome (Barbera *et al*., 2006), and a synthetic LANA peptide with polyethylene glycol and fluorescein was shown to label chromatin in living cells (Fujiwara *et al*., 2021). In the current study, JF646-LANA was synthesized (see Materials and methods) to be used together with sfGFP- and TMR/Cy3-labeled probes. Mitotic chromosomes visualized with Cy3-H3S28ph Fab were indeed more intensely detected by JF646-LANA than by H3K9ac Fab, whose chromatin binding is decreased by deacetylation during mitosis (Fig. S1C and S1D).

To visualize F-actin dynamics at around NEBD, 1-cell stage embryos were injected with UtrCH-sfGFP, TMR-dextran, and JF646-LANA, and were imaged every 12.5 s from 256- cell stage. Figure 4 shows the representative images of cells in the 1k-cell stage (A) and high stage (B) embryos with the changes in N/C intensity ratio of UtrCH-sfGFP and TMR-dextran (C). UtrCH-sfGFP accumulated in nuclei (Fig. 4, −37.5 s to −12.5 s), reaching a peak concomitantly with NEBD as indicated by TMR-dextran diffusing into nuclear areas (Fig. 4, 0 s). After NEBD, UtrCH signals were remained in the vicinity of condensing chromosomes during prometaphase (Fig. 4, 0 s to 25 s), and almost disappeared from the chromosome areas by metaphase (Fig. 4, 37.5 s).

**Figure 4.**
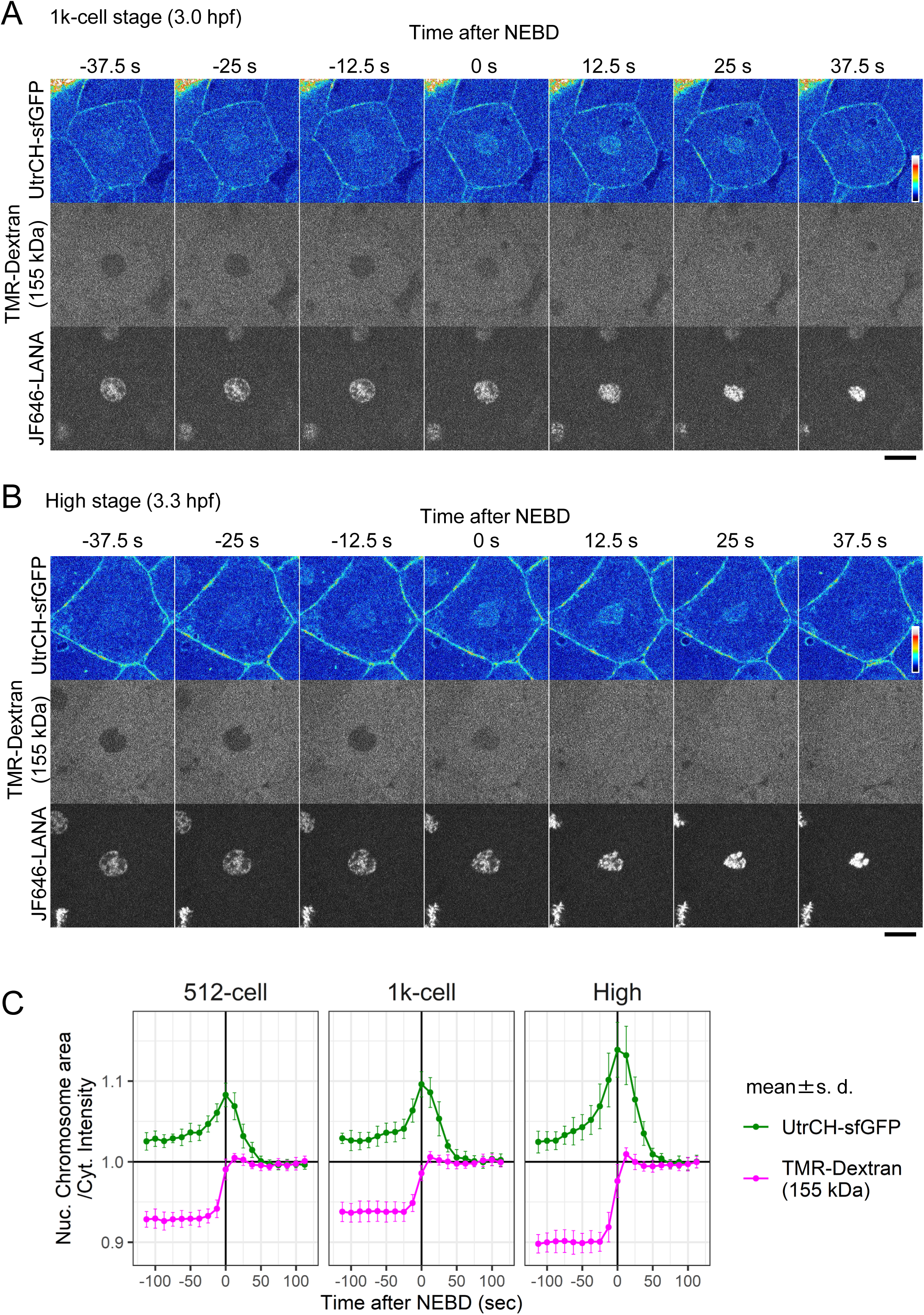
F-actin that accumulated in the nucleus remained there after nuclear envelope breakdown (NEBD). Zebrafish embryos were injected with UtrCH-sfGFP as an F-actin probe, TMR-labeled 155-kDa dextran to monitor NEBD, and JF646-LANA as a chromatin marker. Every 12.5 s, confocal sections were acquired during the 1k-cell (**A**) and the high (**B**) stages. UtrCH- sfGFP accumulated in nuclei (**A** and **B**, −37.5 s to −12.5 s) and remained in the vicinity of chromosomes even after NEBD (**A** and **B**, 0 s to 37.5 s). (**C**) Nucleus/Cytoplasm or Chromosome area/Cytoplasm intensity ratio of UtrCH-sfGFP and TMR-Dextran were measured (N = 10) at the 512-cell, 1k-cell, and high stages. The mean values were shown with the s. d. Nuclear actin levels increased at the late interphase to prophase (from −50 s to 0 s) with a peak at around NEBD (0 s). After NEBD, UtrCH-sfGFP gradually decreased from chromatin (from 0 s to 50 s) during the prometaphase. Scale bars: 20 μm.

To track microtubule dynamics with F-actin and chromosomes during prophase to metaphase, TMR-labeled tubulin was co-injected with UtrCH-sfGFP and JF646-LANA, and three-dimensional images were acquired every 15 s at the 256-cell (Movie 2), 512- cell (Movie 3), 1k-cell (Fig. 5 and Movie 4), high (Fig. 6 and Movie 5), sphere (Fig. 7), and dome stages (Fig. S3). UtrCH-sfGFP was located in the nucleus with enrichment at around the nuclear periphery of the 1k-cell stage embryo before NEBD (Fig. 5, −15 s). Just after NEBD, UtrCH-sfGFP remained in the vicinity of the condensing chromosomes, and microtubule spindles (TMR-Tubulin) grown from centrosomes extended along with the chromosomes (Fig. 5A, 0 s and 15 s; see also Fig. 5B for 3D rendering views). UtrCH- sfGFP signals gradually disappeared from the nuclear areas during prometaphase to metaphase (Fig. 5A, 30–45 s; and Fig. 5B, 30 s). Concomitantly with actin diminishment, microtubule spindles appeared to capture chromosomes for their segregation (Fig. 5, 45 s to 90 s). Similar dynamics of actin and tubulin were observed from the 256-cell stage to 1k-cell stages (256-cell stage: Movie 2, 512 cell-stage: Movie 3). In the oblong stage, the UtrCH-sfGFP signal in the nucleus was much weaker (Fig. 6, −45s to 0 s); however, after the NEBD, the actin patches were still observed in the vicinity of condensing chromosomes (Fig. 6, 15 s and 30 s). In the sphere stage, the timing of UtrCH-sfGFP nuclear accumulation was more transient and the signals were much weaker than those during the 1k-cell stage (Fig. 7, 0 s and 15 s). Only a trace amount of UtrCH-sfGFP was observed in the chromosome area in the dome stage Fig. S3, 30 s and 45 s).

**Figure 5.**
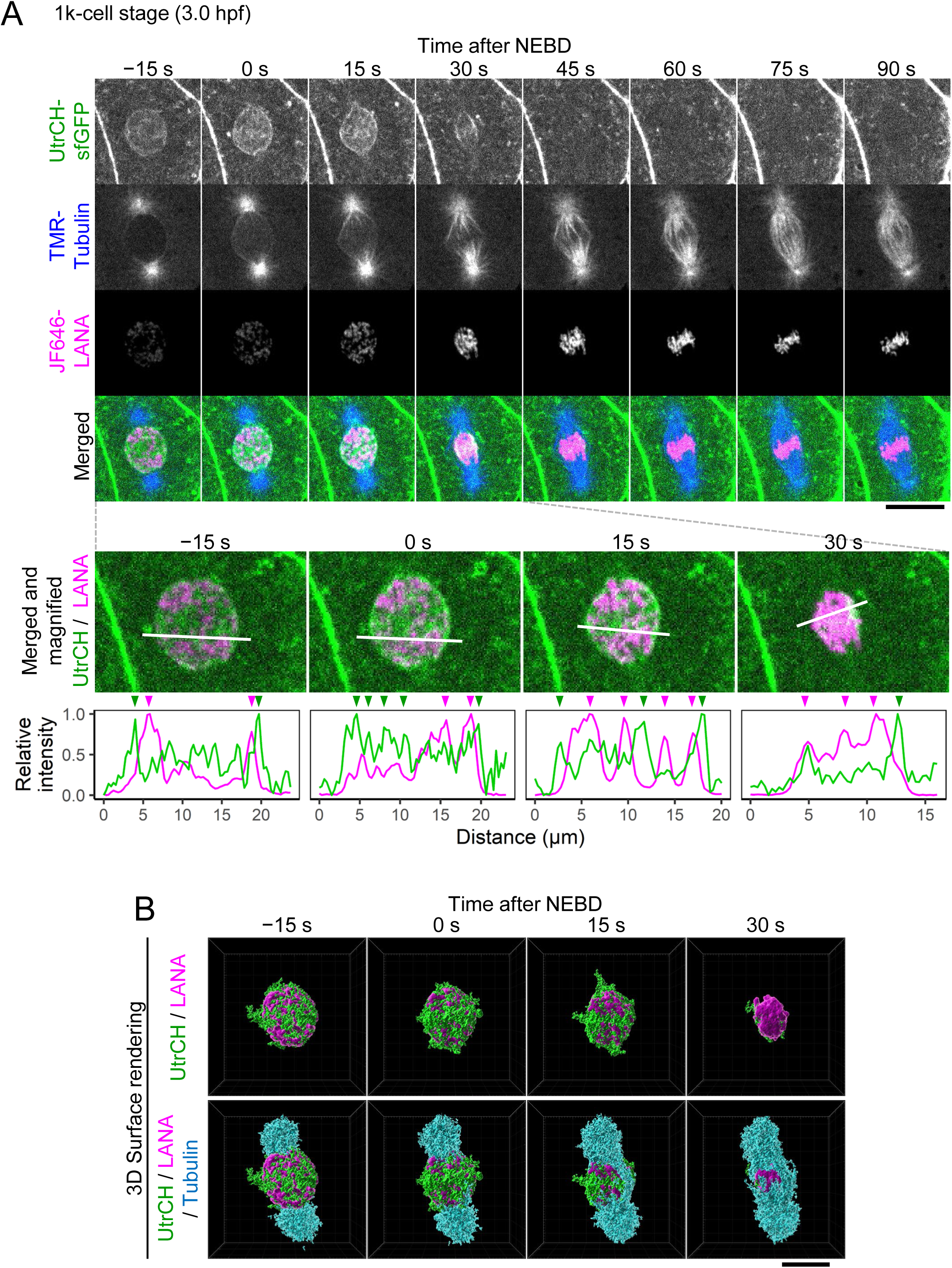
Actin patches remained in the vicinity of chromosomes during prometaphase and disappeared at the metaphase in the 1k-cell stage embryos. Zebrafish embryos were injected with UtrCH-sfGFP, TMR-tubulin, and JF646-LANA. Every 15 s, confocal sections were acquired. (**A**) Single sections of UtrCH-sfGFP, TMR- tubulin, and JF646-LANA at the 1k-cell stage embryo, and their merged and magnified images are shown. Relative intensity profile plots of lines are indicated at the bottom. The intensity is normalized using maximum and minimum intensity. UtrCH-sfGFP signals are located by chromosomes. (**B**) The surface rendering three-dimensional images are shown (green: UtrCH-sfGFP, magenta: JF646-LANA, cyan: TMR-Tubulin). UtrCH-sfGFP accumulated in the nucleus (−15 s) remained in the vicinity of condensing chromosomes after NEBD (from 0 s to 45 s) and disappeared before the chromosomes were aligned at the metaphase (60 s). Just after NEBD (0 s and 15 s), TMR-Tubulin does not appear to reach chromosomes when UtrCH-sfGFP patches are around. When UtrCH-sfGFP patches partially disappear (15 s and 30 s), TMR-Tubulin appears to capture condensing chromosomes (15 s and 30 s). See also Movie 4. Scale bars: 20 μm. See Figure S3 for the sphere stage.

**Figure 6.**
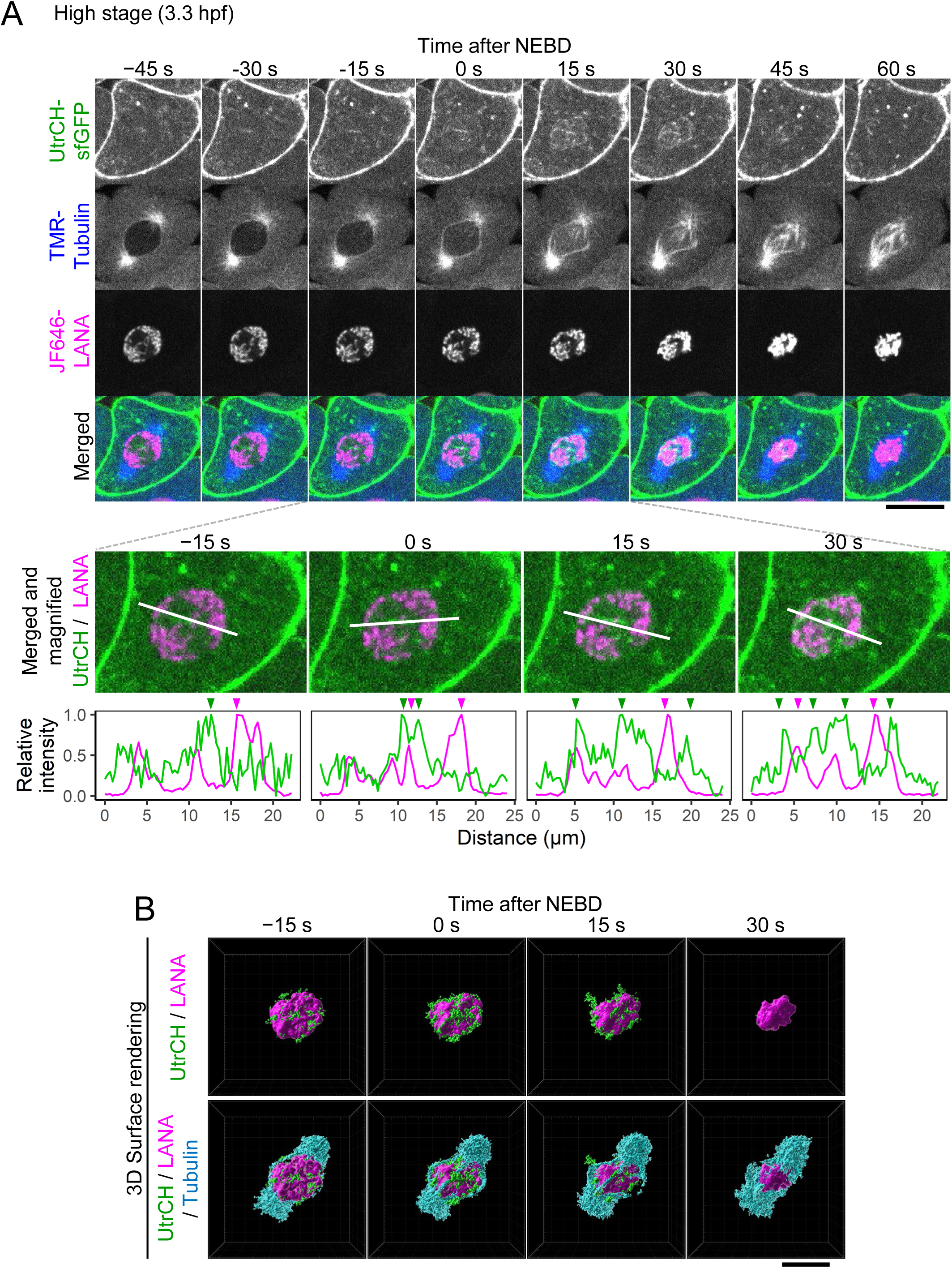
Actin patches partially remained in the vicinity of chromosomes during prometaphase and disappeared at the metaphase in the high stage embryos. Zebrafish embryos were injected with UtrCH-sfGFP, TMR-tubulin, and JF646-LANA. Every 15 s, confocal sections were acquired. (**A**) Single sections of UtrCH-sfGFP, TMR- tubulin, and JF646-LANA at the high stage embryo, and their merged and magnified images are shown. Relative intensity profile plots of lines are indicated at the bottom. The intensity is normalized using maximum and minimum intensity. UtrCH-sfGFP signals are located in the vicinity of chromosomes. (**B**) The surface rendering three-dimensional images are shown (green: UtrCH-sfGFP, magenta: JF646-LANA, cyan: TMR-Tubulin). UtrCH-sfGFP signals appeared around condensing chromosomes after NEBD (from 0 s to 45 s) and disappeared before the chromosomes were aligned at the metaphase (60 s). Just after NEBD (0 s and 15 s), TMR-Tubulin does not appear to reach chromosomes associated with UtrCH-sfGFP patches. TMR-Tubulin appears to capture condensing chromosomes (0 s to 30 s). See also Movie 5. Scale bars: 20 μm.

**Figure 7.**
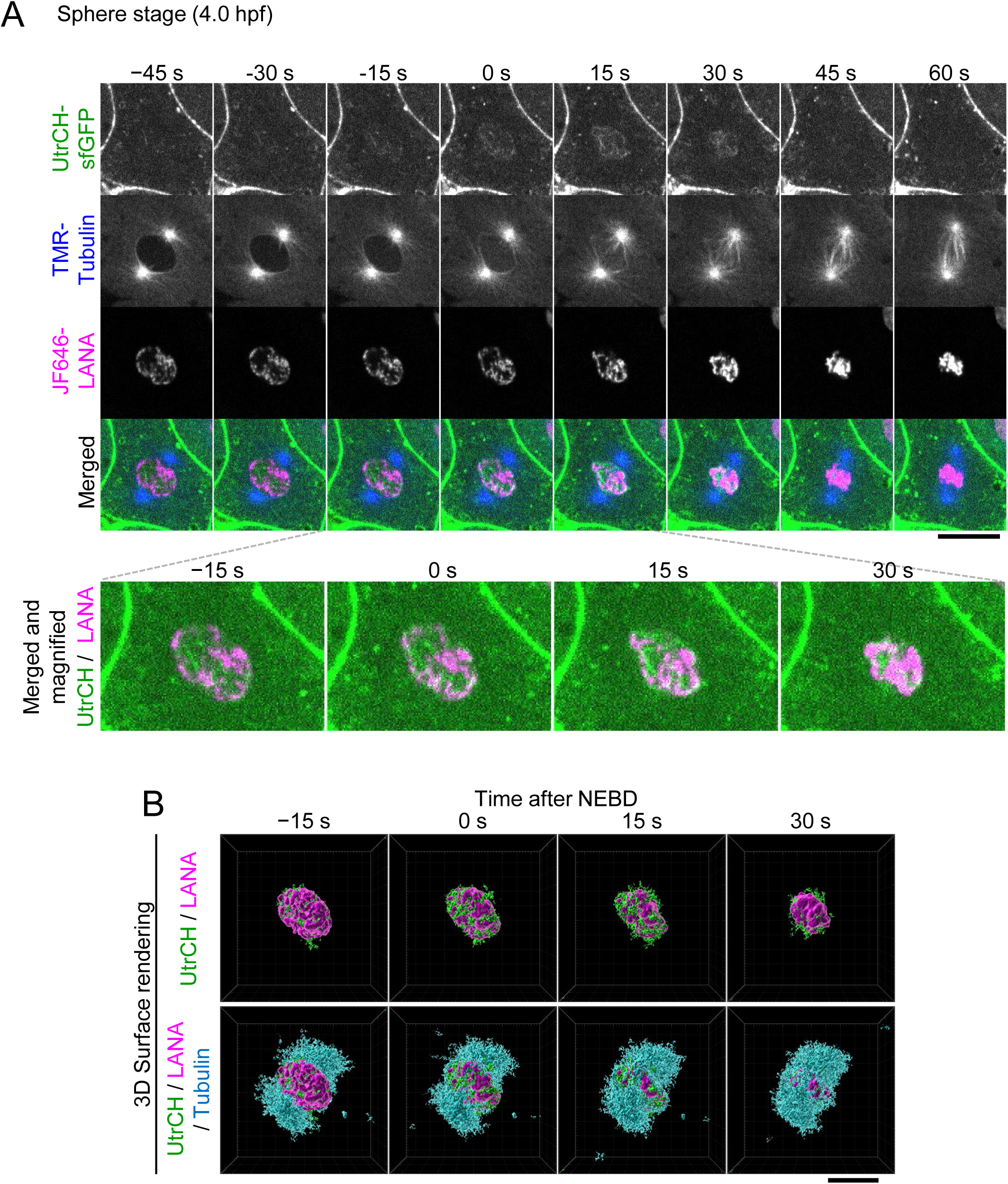
Actin patches appeared during prophase to prometaphase and disappeared at the metaphase in the sphere stage embryo. Zebrafish embryos were injected with UtrCH-sfGFP, TMR-labeled tubulin, and JF646- LANA. Every 15 s, confocal sections were acquired. (**A**) Single sections for UtrCH-sfGFP, TMR-labeled tubulin, and JF646-LANA in sphere stage embryo, and their merged and magnified images are shown. (**B**) The surface rendering three-dimensional images are shown (green: UtrCH-sfGFP, magenta: JF646-LANA, cyan: TMR-Tubulin). Weak UtrCH- sfGFP signals were observed in the vicinity of chromosomes after NEBD (0 s and 15 s) and disappeared (45 s) before chromosomes aligned at the metaphase (60 s). Scale bars: 20 μm.

### Transcription inhibition prolonged nuclear actin decrease after zygotic genome activation (ZGA)

We next examined the relationship between zygotic transcription and nuclear F-actin, because the amount of F-actin was substantially decreased after the oblong stage, as in most somatic cells (Schoenenberger *et al*., 2005; Hofmann *et al*., 2009; Xu *et al*., 2010; Parisis *et al*., 2017). To test whether zygotic transcription is required to diminish nuclear F-actin, we injected α-amanitin, which directly binds to RNA polymerase II and interferes with its translocation and nucleotide incorporation (Nguyen *et al*., 1996; Brueckner and Cramer, 2008), into zebrafish embryos (Fig. 8A; Kane *et al*., 1996; Sato *et al*., 2019). Embryos that were injected with α-amanitin or the buffer were fixed at the 1k-cell and oblong stages and stained with Hoechst and phalloidin. In 1k-cell stage embryos, in which major ZGA typically begins, phalloidin signals were concentrated in the nuclei, regardless of α-amanitin injection (Fig.8B, 1k-cell). In contrast, at the oblong stage, nuclear phalloidin signals were slightly enriched in α-amanitin-injected embryos compared to those detected in the control embryos (Fig. 8B, oblong). We then confirmed the effect of α-amanitin on nuclear F-actin in living cells by using UtrCH-sfGFP. In the both amanitin-treated and control embryos, the N/C ratio of UtrCH-sfGFP began to decrease after the high stage (Fig. 8C). However, compared to the control, the N/C ratio in α-amanitin-injected embryos remained relatively high at sphere and dome stages (Fig. 8C). These data suggest that the decrease of the N/C ratio of UtrCH-sfGFP is facilitated by zygotic transcription.

**Figure 8.**
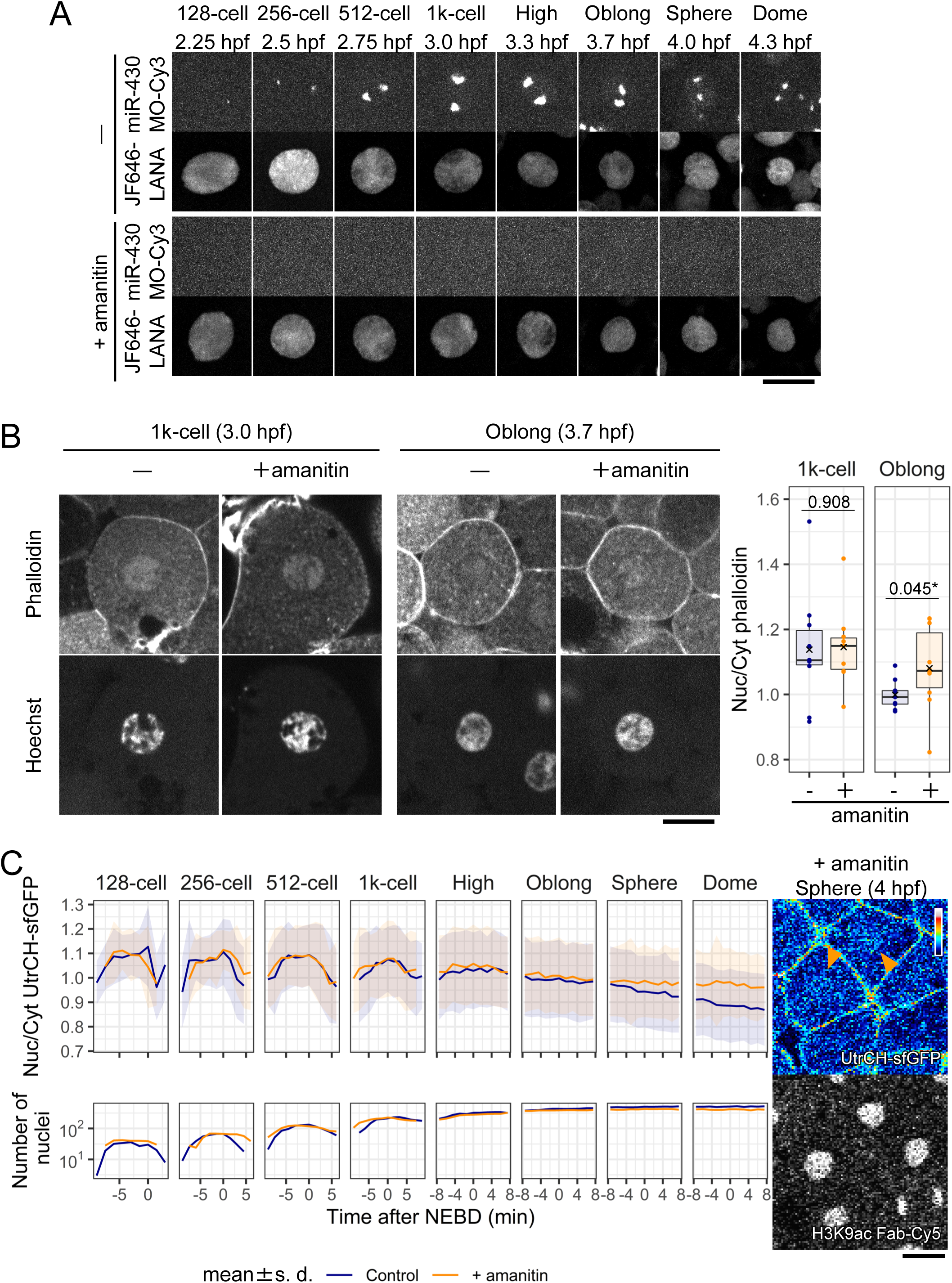
Transcription inhibition delayed the nuclear actin decrease after the 1k- cell stage. (**A**) Zebrafish embryos were injected with α-amanitin to inhibit RNA-polymerase II- mediated transcription, or the vehicle (PBS), with Cy3-labeled morpholino antisense oligonucleotide for miR-430 transcripts as a transcription marker, and JF646-LANA as a chromatin marker. Every 90 s, 25 z-stack images with 4 μm intervals were acquired using a confocal microscope. 7 slices at each time point from the 128-cell to dome stage were depicted and the max intensity projection images are shown. (**B**) Embryos were grown to the 1k-cell stage and oblong stage before fixation and staining with Hoechst and Acti- stain™ 555 phalloidin. Single confocal sections are shown with N/C ratios of phalloidin intensity (N = 10). In the box plots, center lines show the medians; box limits indicate the 25th and 75th percentiles; whiskers extend 1.5 times the interquartile range from the 25th to 75th percentiles; × indicates the means; and data points are plotted as closed circles (PBS, blue; α-amanitin, orange). *p*-values obtained with a Students *t*-test (unpaired, two-tailed) are also shown (*, *p* < 0.05). At the 1k-cell stage, clear phalloidin signals are observed in the nuclei of both embryos treated without or with α-amanitin. At the oblong stage, phalloidin signals are not concentrated in the nucleus of embryos without α-amanitin but are still observed in the nucleus of α-amanitin-treated embryos. (**C**) Zebrafish embryos were injected with α-amanitin, or PBS, and then with UtrCH- sfGFP and H3K9ac Fab-Cy5. Every 90 s, 25 z-stack fluorescence images with 4 μm intervals were acquired. The mean values of N/C ratios (control: blue, +α-amanitin: orange) are plotted with the s. d. from the 128-cell to dome stage. α-amanitin-injected embryos showed the higher N/C ratio of UtrCH-sfGFP value in later stages (sphere and dome). Single confocal sections at the sphere stage are shown on the right. UtrCH- sfGFP remained accumulated in some nuclei during the sphere stage (orange arrowheads). Scale bars: 10 μm (**A**) and 20 μm (**B** and **C**).

### Attempts to disturb nuclear actin in zebrafish early embryos

To investigate the physiological role of nuclear F-actin in early embryos, we sought to artificially manipulate its levels. The first attempt to decrease nuclear actin by ectopic expression of Exp6, which is known to export actin from the nucleus to the cytoplasm (Stüven *et al*., 2003), by injecting mRNA encoding 3×FLAG-tagged zebrafish Exp6 into embryos. However, Exp6-3×FLAG expression induced severe cytokinesis defects but did not decrease nuclear actin levels (Fig. S4).

Next, we expressed a polymerization-deficient actin mutant (R62D), in which Arg 62 was substituted to Asp (Posern *et al*., 2002), tagged with the superfolder Cherry and a nuclear localizing signal (Actin R62D-sfCherry-NLS), to inhibit nuclear F-actin formation. The wild type actin tagged with sfCherry-NLS (Actin WT-sfCherry-NLS) and sfCherry-NLS were used as controls to examine the effect of the wild-type actin and sfCherry alone. Embryos were co-injected with mRNA encoding Actin R62D-sfCherry-NLS, Actin WT-sfCherry-NLS, or sfCherry-NLS, with UtrCH-sfGFP and Cy5-labeled H3K9ac- specific Fab and were imaged from 8-cell stage. The N/C intensity ratios of UtrCH-sfGFP, however, did not show obvious differences among embryos expressing sfCherry-NLS, Actin R62D-sfCherry-NLS, and Actin WT-sfCherry-NLS (Fig. S5A and S5B). Expression of Actin R62D tagged with smaller 3xFLAG tagged versions, instead of sfCherry, also did not affect the N/C ratios of UtrCH-sfGFP (Fig. S5C). Because actin was already abundantly present in nuclei, additional expression of the wild-type or mutant actin might have minimal effects.

In has been reported that cofilin proteins actively depolymerize actin filaments in cytoplasm during the interphase (Vartiainen *et al*., 2002; Amano *et al*., 2002; Kaji *et al*., 2008; Chin *et al*., 2016; Wioland *et al*., 2017). Cofilin1 is the non-muscle isoform expressed ubiquitously while cofilin2 is the muscle isoform. We expressed zebrafish cofilin1 and cofilin2, tagged with sfCherry and an NLS, to attempt depolymerizing nuclear actin in embryos, but again the N/C intensity ratios of UtrCH-sfGFP were not affected (Fig. S6). Thus, it was not possible to manipulate the level of nuclear F-actin in our experimental conditions. Nuclear F-actin level might be robustly regulated in zebrafish early embryos.

## Discussion

### Nuclear actin accumulation in zebrafish early embryos

We demonstrated F-actin accumulation in zebrafish early embryo nuclei by fixed cells using phalloidin and by living cells using fluorescent actin-binding probes. In living embryos from the 8-cell to 1k-cell stages, UtrCH-sfGFP, which selectively binds to F-actin, gradually accumulated in the nucleus during the interphase, thereby reaching the maximum amount at around NEBD. After the high stage, UtrCH-sfGFP was not much concentrated in nuclei, while some filamentous structures were more evidently observed. These data are consistent with previous findings in mouse and *Xenopus laevis* early embryos (Baarlink *et al*., 2017; Oda *et al*., 2017), which indicates that nuclear F-actin accumulation in undifferentiated cells is common in vertebrates (Xu *et al*., 2010; Miyamoto *et al*., 2011; Miyamoto *et al*., 2013; Sokolova *et al*., 2018; Okuno *et al*., 2020). After ZGA, the level of nuclear F-actin drops. The decrease appears to be induced by zygotic transcription, as nuclear F-actin was still observed in the sphere-stage embryos that were treated with α-amanitin, an RNA polymerase II inhibitor. Maternal-to-zygotic transition may alter the balance of nuclear import and export factors of actin, as it was previously demonstrated that the actin exporter, Exp6, is not expressed in *Xenopus* oocytes nor in early embryonic stages, yet it becomes expressed after ZGA (Bohnsack *et al*., 2006).

### Nuclear F-actin may facilitate chromosome congression and regulate microtubule growth

In early-stage zebrafish embryos, the peak of nuclear F-actin accumulation was concurrent with the timing of NEBD and F-actin localized in the vicinity of condensing chromosomes during prometaphase before disappearing during metaphase. This actin behavior is similar to that observed in starfish oocytes (Lénárt *et al*., 2005; Mori *et al*., 2011; Mori *et al*., 2014; Wesolowska *et al*., 2020). In starfish oocyte GV, F-actin is assembled underneath the nuclear membrane and forms spikes toward to the nuclear membranes, which contribute to membrane rupture and NEBD (Mori *et al*., 2014; Wesolowska *et al*., 2020). After the germinal vesicle breakdown, actin meshwork catches chromosomes and shrinks like a fishnet to transport them over a long distance toward underneath the starfish oocyte cortex in meiosis II (Lénárt *et al*., 2005; Mori *et al*., 2011). In the mouse oocyte, both nuclear F-actin and spindle F- actin protect the chromosomes misalignment and missegregation at the meiosis (Mogessie and Schuh, 2017; Scheffler et al., 2022). It is interesting to speculate that the function of actin in large oocyte cells may partially be retained in fertilized eggs, although it is unknown whether a similar system to starfish is conserved in zebrafish oocytes. During the short cell cycle of zebrafish embryos, the spindle checkpoint is weak until the zygotic genome activation (Ikegami *et al*., 1997). In relatively large cells during the early embryonic stages, actin patches remained around chromosomes may assist in chromosome congression, just as the actin meshwork in starfish oocytes. It is also possible that actin patches guide microtubule assembly and promote proper and rapid spindle formation, as recent reports have shown that actin supports proper spindle formation in *Xenopus* gastrula embryos, mouse and human oocytes, and cultured cells (Woolner *et al*., 2008; Mogessie and Schuh, 2017; Roeles and Tsiavaliaris, 2019; Plessner *et al*., 2019; Farina *et al*., 2016; Farina *et al*., 2019; Kita *et al*., 2019). However, “spindle-actin” has been depicted with barrel structures composed of fine actin fiber at metaphase, unlike actin patches during the prometaphase of zebrafish embryos. We speculate that the accumulation of F-actin in nuclei in zebrafish early embryos contribute to proper mitotic progression by assisting in NEBD, chromosome congression, and/or spindle assembly under rapid cell cycles with weak checkpoint mechanism.

### Manipulation of nuclear actin level in zebrafish embryos

We took different approaches to decrease the level of nuclear F-actin by expressing Exp6, an NLS-tagged Actin R62D, and NLS-tagged cofilin; unfortunately, however, none of them were successful. Exp 6 and NLS-tagged Actin R62D have been used to decrease nuclear actin and inhibition of nuclear actin polymerization, respectively, in previous reports (Bohnsack *et al*., 2006; Okuno *et al*., 2020; Miyamoto *et al*., 2011; Spencer *et al*., 2011; Dopie *et al*., 2012; Parisis *et al*., 2017; Baarlink *et al*., 2017). However, we did not observe significant decreases of UtrCH-sfGFP N/C ratios in zebrafish embryos expressing Exp6 and Actin R62D, under the conditions that did not affect cytoplasmic F-actin. Increasing the expression levels of those proteins by injecting more mRNA affected cytoplasmic F-actin causing defects in cytokinesis. We also attempted to decrease nuclear F-actin using NLS-tagged cofilin 1 and cofilin 2, which are actin-depolymerizing factors (Amano *et al*., 2002; Vartiainen *et al*., 2002; Kaji *et al*., 2008; Chin *et al*., 2016; Wioland *et al*., 2017); however, their expression did not affect nuclear F-actin levels. In addition, the expression of NLS-tagged wild-type actin did not increase UtrCH-sfGFP N/C ratio. It can be speculated that nuclear actin is very abundant and the ectopic wild-type actin or R62D mutant may not affect much on the total F-actin level. Also, F-actin formation in the nucleus may be robustly regulated, and expressing factors that assist in actin depolymerization may be compensated when the expression level is mild not affecting the cytoplasmic actin.

## Materials and methods

### Zebrafish embryo preparation for live imaging

To obtain fertilized eggs, 5–10 pairs of AB line WT zebrafish (*Danio rerio*) males and females were crossed. Embryos were dechorionated using 1 mg/mL pronase from *Streptomyces griseus* (SIGMA, 10165921001) diluted in 0.3× Danieau’s buffer [17.4 mM NaCl, 210 μM KCl, 120 μM MgSO4, 180 μM Ca(NO3)2, 1.5 mM HEPES, pH 7.6]. For actin visualization, UtrCH-sfGFP (1.46 fmol; see below) or LifeAct-sfGFP (2.3 fmol; see below) was injected with labeled Fabs (anti-H3K9ac [CMA315; 50 pg], anti- H3S28ph [CMA315; 50 pg]; anti-DYKDDDDK tag [FUJIFILM Wako, 012-22384; 50 pg]; Sato *et al*., 2019) and 10 fmol JF646-LANA (see below) in 0.5 nL phosphate-buffered saline (PBS, Takara, T900) containing 0.05% phenol red (SIGMA, P5530) as an injection marker, were injected into the yolk of 1-cell stage embryos. For NEBD or tubulin visualization, TMR-labeled 155-kDa dextran (250 pg in 0.5 nL PBS, SIGMA, T1287) or TMR-labeled tubulin (1.25 ng in 0.5 nL General Tubulin Buffer, Cytoskeleton, TL590M) was injected with UtrCH-sfGFP and JF646 LANA. When mRNA was also injected, mRNA of sfCherry-NLS or 3×FLAG-tagged Actin WT or R62D (125 pg in 0.5 nL water), 3×FLAG-tagged Exp6 (150 pg in 0.5 nL water), sfCherry-NLS-tagged cofilin1 (125 or 400 pg in 0.5 nL water) or sfCherry-NLS-tagged cofilin2 (125 pg in 0.5 nL water) was first injected into the cell of 1-cell stage embryos, and then UtrCH-sfGFP, LifeAct-sfGFP, Fabs, and/or JF646-LANA were injected into the yolk. For transcription inhibition, 250 pg α-amanitin (Merck, 129741) in 0.5 nL PBS was injected into the yolk of 1-cell stage embryos (Kane *et al*., 1996; Sato *et al*., 2019). To monitor the transcription inhibition, 530 pg of Cy3- labeled antisense morpholino oligonucleotide specific to miR-430 transcripts (Hadzhiev *et al*., 2019) in 0.5 nL water was injected into the yolk of the 1-cell stage embryos with JF646-LANA. The injected embryos were incubated at ∼25°C up to the 4-cell stage and embedded into 0.5% agarose type VII-A low melting temperature (SIGMA, A-0701) in PBS on a glass bottom dish (IWAKI, 3791-035) with the animal pole positioned down for live-cell imaging.

All zebrafish experiments were approved by the Tokyo Institute of Technology Genetic Experiment Safety Committee (I2018001), and animal handling was carried out according to the guidelines.

### Zebrafish embryo fixation and staining

For Fig. 1A and B, dechorionated embryos were incubated until fixation at the desired stage with 4% paraformaldehyde in 250 mM HEPES for 3 h at 25°C. After fixation, embryos were washed three times with PBS and stained with 2 μg/mL Hoechst 33342 (Nacalai Tesque, 04929-82) and 20 nM Acti-stain™ 555 phalloidin (Cytoskeleton, PHDH1-A) in PBS containing 0.05% Triton X-100 at 4°C overnight in the dark. Stained embryos were washed three times with PBS before microscopic observation. For Fig. 8B, embryos were injected with α-amanitin (250 pg in 0.5 nL) or PBS (0.5 nL) as the control and incubated until the 1k-cell or oblong stage and fixed with 4% paraformaldehyde in 250 mM HEPES for 3 h at 25°C. After the fixation, embryos were washed with PBS and embedded in 100 μL 0.5% agarose in PBS and covered with 100 μL 1% agarose to prevent the detachment of embryos from a glass bottom dish. Embryos were stained with 2 μg/mL Hoechst 33342 and 20 nM Acti-stain™ 555 phalloidin in PBS containing 0.05% Triton X-100 in the dark with gentle shaking (Titec, Shake-XR; 40 rpm) overnight. Stained embryos were washed three times with PBS before microscopic observation.

### Embryo imaging

Images were collected using a confocal microscope. For Figs. 1, 2, 3, 8A, 8C, S2, S4, S5, S6, and Movie 1, images were acquired using a confocal microscope FV1000 (OLYMPUS) operated under built-in software (FLUOVIEW-ASW, version 4.2) with a UPLSAPO 30× silicone-immersion objective lens (NA 1.05), with 150 μm pinhole, zoom 1.0×, and 488-nm, 543-nm, and 633-nm laser lines on a heated stage at 28°C (Tokai Hit). For live imaging of Figs. 1D, 2A, 3A, 8C, S2, S4, S5, S6, and Movie 1, 25 z-stack images (4 μm z-intervals; 512 × 512 pixels) were taken with 90 s intervals.

For high-speed or higher resolution imaging to produce the images shown in Figs. 4, 5, 6, 7, 8A, S1C, S1D, S3, and Movie 2–5, images were taken using a spinning disk confocal system consisting of an inverted microscope (Ti-E, Nikon) with an Apo 40× water-immersion objective lens (NA 1.25; Figs. 4, 5, 6, 7, 8A, S3 and Movies 2–5) or a Plan Apo 25× silicon-immersion objective lens (NA 1.05, Fig. S1C and S1D), a spinning disk unit with 40 μm pinhole (Dragonfly, Oxford Instruments), a laser unit with 488-nm, 561-nm and, 637-nm laser lines, and an EM-CCD camera (iXon Life 888, Oxford Instruments), operated under Fusion software (version 2.0 and 2.2, Oxford Instruments). 15–25 z-stack images (1, 2, or 4 μm z-intervals; 1024 × 1024 pixels) were taken with 12.5, 15, 60, or 90 s time intervals at 25°C.

### Actin-binding peptides

DNA fragment encoding UtrCH (1-262 of human utrophin, NM_007124) or LifeAct (Riedl *et al*., 2008) gifted from Dr. Mari Iwabuchi (Nagoya University, Japan) was fused with the sfGFP gene with a linker sequence (DPPVAT) and was cloned into a pGEX6P-1 vector that has the N-terminal glutathione S-transferase (GST) tag (Cytiva). *E. coli* BL21 (DE3) cells harboring the pGEX6P-1-based expression plasmid were grown in 24 mL Plusgrow II (Nacalai Tesque, 08246-86) medium overnight at 25°C. The overnight culture (24 mL) was diluted into 400 mL PlusgrowII medium and further incubated at 25°C for 3 h. The expression of GST-UtrCH-sfGFP and GST-LifeAct-sfGFP was induced in *E. coli* BL21 (DE3) at 25°C for 7 h by adding 1 mM isopropyl β-D-thiogalactopyranoside (Nacalai Tesque, 19742-94). Harvested bacterial cells (400 mL, 4,000 ×*g*, 10 min, 4°C) were suspended in 20 mL ice-cold 50 mM Tris–HCl (pH 7.4) buffer containing 1 mg/mL lysozyme (Seikagaku Corporation, 100940) and 1× protease inhibitor cocktail (Nacalai Tesque, 03969-21) and were lysed by sonication (Branson; SONIFIRE 250; output control: 1, duty cycle: 50% for 2 min repeated 14 cycles with 30 s interval) in an ice bath. Triton X-100 (10% in water) was added to the sonicated solution to yield the final concentration at 1% and the mixture was incubated for 30 min at room temperature with a gentle rotation (Titec, RT-50; 30 rpm). After centrifugation (10,000 ×*g* for 20 min at 4°C), the supernatant was mixed with 5 mL (bed volume) glutathione sepharose 4B beads (Cytiva, 17075605; prewashed with PBS) and gently rotated (Titec, RT-50; 30 rpm) at 4°C for 20 h in the dark. The beads were collected in a Poly-Prep® Chromatography Columns (BioRad) and washed with a 10-column volume of PBS. GST-UtrCH-sfGFP or GST-LifeAct-sfGFP was eluted using a 5-column volume of 10 mM glutathione (FUJIFILM Wako Pure Chemical, 073-02013) containing 50 mM Tris–HCl (pH 8.7). The eluate was concentrated down to 1.5 mL using an Amicon Ultra Centrifuge Filter unit (30 kDa cutoff for GST-UtrCH-sfGFP and 10 kDa cutoff for GST-LifeAct-sfGFP; Merck, UFC903024 and UFC901024, respectively). The GST tag was removed by digestion with 8 units/mL PreScission Protease (Cytiva, 27084301) on ice for 24–36 h and then removed through glutathione sepharose 4B beads (5 mL bed volume) packed twice in Poly-Prep® Chromatography Columns (BioRad). UtrCH-sfGFP and LifeAct-sfGFP were further purified through gel filtration column chromatography (HiLoad 16/600 Superdex 75 pg, Cytiva, 28989333) using AKTAPrime™PLUS (Cytiva, 11001313). After loading 1.5 ml of the protein sample, 30 mL fractions were collected and analyzed by 15% sodium dodecyl sulfate (SDS)-polyacrylamide gel electrophoresis (SuperSep Ace, FUJIFILM Wako Pure Chemical, 190-15001). The highly pure fractions were collected (Supplementary Fig. S1A and S1B), aliquoted, snap-frozen in liquid N2, and stored at −80°C. Once thawed, recombinant proteins were stable at 4°C for at least 5 days.

### Fab preparation

Dye-conjugated Fabs were prepared as previously described (Kimura and Yamagata, 2015). In brief, 100 μg purified Fab (anti-H3K9ac: CMA310/19E5, Hayashi-Takanaka *et al*., 2011; anti-H3S28ph: CMA315/10-20F11, Hayashi-Takanaka *et al*., 2014; anti-DYKDDDDK tag: FUJIFILM Wako, 012-22384) was reacted with N-hydroxysuccinimide ester-conjugated fluorescence dye (Cy5, cytiva, PF11A25001; Cy3, cytiva, PA13105) in 100 mM NaHCO3 (pH 8.3) in PBS for 1 h at room temperature in the dark. Dye-conjugated Fabs were collected using PD MiniTrap G-25 column (Cytiva, 28918004; pre-equilibrated with PBS) and were concentrated to 1 mg/mL using a 10-kDa cutoff Amicon Ultracell Centrifuge Filter Unit (Merck, UFC5010BK), and stored at 4°C in the dark.

### Morpholino oligonucleotide labeling

Antisense morpholino oligo nucleotide was labeled as previously described (Sato *et al*., 2019). In brief, 3′-primary amino-modified morpholino oligonucleotides specific for miR- 430 was reacted with N-hydroxysuccinimide ester-conjugated Cy3 in 0.1 M NaHCO3 (pH 8.3) for 1 h at room temperature in the dark. After the reaction, Cy3-labeled morpholino oligonucleotide was purified using a PD MiniTrap G-25 column.

### Synthesis of PEG750-LANA-JF646 (JF646-LANA)

Preparative high-performance liquid chromatography (HPLC) was conducted by using a JASCO HPLC system equipped with a UV-2075 spectrometer (230 nm), PU-4086 pumps, a DG-4580 degasser, and an MX-2080-32 mixer using a YMC-Triart C18 (20 mm I.D. × 250 mm or 10 mm I.D. × 250 mm) column at 40°C with a gradient of acetonitrile in 0.1% aqueous trifluoroacetic acid (TFA) at a flow rate of 10 or 3.5 mL/min. Analytical HPLC was conducted by using a JASCO HPLC system equipped with a UV-2075 spectrometer, PU-4180 pumps, a DG-4580 degasser, and an MX-2080-32 mixer, using a YMC-Triart Phenyl C18 (4.6 mm I.D. × 150 mm) column with a linear gradient of 2%–90% acetonitrile in 0.1% aqueous TFA over 3–15 min.

Poly(ethylene glycol) methyl ethers (Mn = 750) were purchased from Sigma-Aldrich (#202495). All protected α-amino acids were purchased from Watanabe Chemical Industries, Ltd. (Hiroshima, Japan) and Peptide Institute, Inc. (Osaka, Japan). NovaPEG Rink amide resin was purchased from Merck KGaA, Co. Other chemicals were used as received, unless otherwise stated.

The LANA peptide was synthesized on a solid phase using NovaPEG-Rink-amide-resin (Fig. S7A). The Fmoc-amino acid was sequentially coupled using the DIC-HOBt method in N,N-dimethylformamide (DMF; 3 equiv. each) for 60 min at room temperature after removal of each Fmoc group with 20% piperidine-DMF for 10 min. The N-terminus was capped with 20% acetic anhydride in CH2Cl2 for 5 min. The peptide was cleaved from the resin by treatment with TFA in the presence of triisopropylsilane (TIPS) and H2O (95:2.5:2.5) for 90 min at room temperature, and then was concentrated under reduced pressure, and precipitated with ether to obtain crude peptides, which were purified with preparative HPLC (0% acetonitrile for 5 min, followed by a linear gradient of 0%–60% acetonitrile over 30 min with a flow rate of 10 mL/min) to afford the product LANA5-22-N3 peptide (Fig. S7B) as white solids after lyophilization. 3,4,5-Tris(PEG750)-N-(prop-2-yn-1- yl)benzamide was synthesized (Fig. S7C) following a previously described method (Fujiwara *et al*., 2021).

To synthesize PEG750-LANA (Fig. S7D), Cu-Tris[(1-benzyl-1H-1,2,3-triazol-4- yl)methyl]amine (TBTA) solution in water:tBuOH (1:2, 150 μL; separately prepared by mixing 24 mM CuSO4 solution in water and 24 mM TBTA solution in tBuOH) was added to a stirred solution of LANA5-22-N3 (10 mM in water, 72 μL, 720 pmol) and 3,4,5-tris(PEG750)-N-(prop-2-yn-1-yl)benzamide (50 mM in tBuOH, 14.4 μL, 720 pmol). Then, sodium ascorbate aqueous solution (100 mM, 60 μL, 6.00 μmol) was added, and the mixture was stirred at room temperature for 22 h. Insoluble materials were removed by filtration, and the filtrate was purified with preparative HPLC (0% acetonitrile for 3 min, followed by a linear gradient of 0%–25% acetonitrile over 2 min, then a linear gradient of 25%–80% acetonitrile over 35 min with a flow rate of 3.5 mL/min) to produce PEG750- LANA; (Fig. S7E) after lyophilization.

To synthesize PEG750-LANA-JF646 (Fig. S7F), 276 μL of DMF was added to a stirred solution of PEG750-LANA (10 mM in water, 40 μL, 400 pmol) and Janelia Fluor 646, N-succinimidyl ester (TOCRIS, 10 mM in DMSO, 44 μL, 720 pmol). Then, 40 μL saturated sodium hydrogen carbonate aqueous solution was added, and the mixture was stirred at room temperature for 5 h. Insoluble materials were removed by filtration, and the filtrate was purified with preparative HPLC (0% acetonitrile for 3 min, followed by a linear gradient of 0%–25% acetonitrile over 2 min, then a linear gradient of 25%–80% acetonitrile over 35 min with a flow rate of 3.5 mL/min) to produce PEG750-LANA-JF646 (Fig. S7G) after lyophilization. The final product was analyzed by a MALDI-TOFMS (Fig. S8, Shimadzu Biotech Axima ToF^2^ spectrometer).

### mRNA synthesis

To clone zebrafish genes, RNA was prepared from 24 hpf embryos using TRIzol reagent (Ambion, 15596026). Zebrafish WT beta-actin (actin WT, GenBank Accession: AF025305), Exp6 (GenBank Accession: BC044132), cofilin1 (GenBank Accession: AY398323), and cofilin2 (GenBank Accession: AY398324) cDNA were amplified from the embryo RNA using PrimeScript™ II High Fidelity One Step RT-polymerase chain reaction (PCR) Kit (Takara, R026A) using the following primers: actin WT: (5’- ATGGATGATGAAATTGCCGCACTGGTTGT-3’) and (5’- GAAGCATTTGCGGTGGACGATGGATGGTCC-3’); Exp6: (5’- ATGGCGTCAGAGGAGGCCTCCTTACGGGCC-3’) and (5’- TAACTTCAGCGTTCCCGGCGGGAGGCTCCC-3’); cofilin1: (5’- ATGGCCTCAGGTGTAGC-’3) and (5’-TTAGACAGGCTTCCCCTCC-’3); cofilin2 (5’- ATGGCCTCCGGAGTTAC-3’) and (5’-TCAATCGGTTAGAGGCTTTCC-3’). Amplified cDNA was cloned into pSC plasmid (StrataClone Blunt PCR Cloning Kit, STRATAGENE, 240207). Then, the actin R62D mutant was obtained by inverse PCR using the following primers (5’-GATGAGGCTCAGAGCAAGAGAGGTATCCTGACCCTGAAG-3’) and (5’- CTTCAGGGTCAGGATACCTCTCTTGCTCTGAGCCTCATC-3’). The nucleotide sequence was validated by Sanger sequencing. For *in vitro* transcription, actin WT and R62D were inserted into the EcoRI and NotI sites of the pcDNA3.1 + poly(A)83 plasmid (Yamagata *et al*., 2005) with sfCherry or 3×FLAG-tag and NLS (see below). Exp6 was inserted into the EcoRI and NotI sites of the pcDNA3.1 + poly(A)83 plasmid with 3×FLAG-tag. Cofilin1 or cofilin2 were inserted into the EcoRI and NotI sites of the pcDNA3.1 + poly(A)83 plasmid with sfCherry and NLS. The resulting pcDNA3.1_sfCherry-NLS + poly(A)83, pcDNA3.1_actin R62D-sfCherry-NLS + poly(A)83, pcDNA3.1_actin WT-sfCherry-NLS + poly(A)83, pcDNA3.1_actin R62D-3×FLAG-NLS + poly(A)83, pcDNA3.1_actin WT-3×FLAG-NLS + poly(A)83 pcDNA3.1_sfCherry-NLS + poly(A)83, pcDNA3.1_Cofilin1-sfCherry-NLS + poly(A)83, pcDNA3.1_Cofilin2-sfCherry- NLS + poly(A)83 were linearized by XbaI digestion, and pcDNA3.1_Exp6-3×FLAG-NLS + poly(A)83 was linearized by XhoI digestion. The linearized plasmids were treated with 0.2 mg/mL proteinase K (Invitrogen, 25530-015), purified with phenol–chloroform treatment and ethanol precipitation, then processed through *in vitro* transcription using mMESSAGE mMACHINE T7 Transcription Kit (Ambion, AM1344) according to the manufacturer’s instruction. The concentration of synthesized mRNA was measured and adjusted to 50, 250, 300, or 800 ng/μL by using nuclease-free water (Cytiva, SH30538.01) containing 0.05% phenol red (SIGMA-ALDRICH, P5530-5G) as an injection marker.

### Nuclear Localization Signal cloning

Nucleoplasmin 2B (NPM, GenBank Accession: NM_001123007) was amplified from 24 hpf embryo RNA using the PrimeScript II High Fidelity One Step RT-PCR kit (Takara, R026A) and the following primers; 5’-ATGAGCAAAACCGAGAAACC-3’ and 5’- TCACGCCTTCGCCTTC-3’. The nucleotide sequence was validated with Sanger sequencing. NPM cDNA was fused to the C-terminus of sfGFP and cloned into pcDNA3.1 containing a poly(A) tail sequence for mRNA synthesis, as described above. After confirming the nuclear localization of sfGFP-NPM, the potential NLS region containing 41 amino acids at the C-terminus was subcloned into the same vector. The 41 amino acids directed sfGFP to the nuclei, so were used as an NLS in zebrafish embryos.

### Western blotting

After dechorionation and injection, zebrafish embryos were incubated at 28°C for 5.3 h. At the 50%-epiboly stage, the yolk was removed by hand using a pair of sharp forceps and the cells were washed three times in 0.3× Danieau’s buffer. Ten de-yolked embryos were transferred into a 1.5 ml protein low bind-tube (Eppendorf, 0030108116) with the buffer and the number of the embryos was confirmed under a stereoscopic microscope.

A gel-loading tip (QSP, 010-R204S-Q) was used to remove as much of the extra buffer as possible. Then, 10 μL 2× SDS-gel loading buffer (125 mM Tris–HCl, 0.01% bromophenol blue, 4% SDS, 20% glycerol, 100 mM DTT) was added to the embryos, mixed by vortex and boiled at 95°C for 5 min. The 10 embryos were loaded into each lane and separated on a 7.5% polyacrylamide gel (SuperSep Ace, FUJIFILM Wako Pure Chemical, 191-14931), then transferred onto a polyvinylidene fluoride membrane (PALL, BSP0161) using a semi-dry blotting system (ATTO). After blocking with Blocking-One (Nacalai Tesque, 0395395), the membrane was incubated with anti-DYKDDDDK antibody (1E6, 1 μg/mL, FUJIFILM Wako Pure Chemical, 012-22384) overnight at 4°C. After washing with TBST (0.01% Tween-20, 10 mM Tris, 150 mM NaCl, pH 8.0), the membrane was incubated with horseradish peroxidase-conjugated goat anti-Mouse IgG (1:1000 dilution, Jackson ImmunoResearch, 115-035-146) overnight at 4°C. After washing the membrane with TBST, the signal was developed using Western Lightning® Plus-ECL (PerkinElmer, NEL104001EA) and detected using a gel documentation system (ATTO, LuminoGraphII).

### Data analysis

The intensity ratio of the nucleus to the cytoplasm depicted in Figs. 2, 3, 8C, S4, S5, and S6 were measured as previously described (Sato *et al*., 2019) using MATLAB software (https://github.com/lhilbert/NucCyto_Ratio_TimeLapse). Using this code, the nuclear area was segmented by the Cy5-labeled H3K9ac channel, and the cytoplasm was defined as the region between 3.3 μm and 8.3 μm from the surface of the nucleus. To measure the intensity ratio of the nucleus or chromosomes to the cytoplasm of UtrCH shown in Fig.4C and 8B, a single z-section of the equatorial plane of each nucleus was selected, and the nuclear area was determined based on the JF646- LANA or Hoechst signal. The cytoplasmic area was defined as the region between 0.3 μm and 2.5 μm from the edge of the nucleus using ImageJ FIJI (ver. 2.1.0).

Drawing line graphs, box plots, and statistical analyses were performed by using R software (version 4.0.3; https://www.r-project.org/). The data distribution of each dataset was first analyzed using Kolmogorov-Smirnov test. For parametric samples, the homoscedasticity was analyzed using *F*-test, and then Student’s *t*-test (unpaired, two- tailed, Fig. 7A; comparing N/C ratio with homoscedasticity) was used.

## Supporting information

8-cell to high st.

256-cell st.

512-cell st.

1k-cell st.

High st.

## Acknowledgement

We appreciate Mari Iwabuchi and Kazuo Yamagata for plasmids, Lennart Hilbert for MatLab code, and Masahiko Harata, Kei Miyamoto and members in Kimura lab for discussion. We thank the Biomaterials Analysis Division, Open Facility Center, Tokyo Institute of Technology for DNA sequencing.

