## Supplementary information for "Actin filaments accumulate in the nucleus during interphase and remain in the vicinity of condensing chromosomes during prophase to metaphase in zebrafish early embryos"

**Supplementary figures**

Figure S1. Actin and chromatin probes.

Figure S2. Actin accumulated in the interphase nuclei at the 16-, 64-, 256-, and 1k-cell stages.

Figure S3. Weak actin patches in the dome stage embryo.

Figure S4. Expotin 6 expression did not decrease nuclear actin levels but caused severe defects in cytokinesis.

Figure S5. NLS-tagged polymerization-defective mutant and wild-type actin proteins did not alter nuclear F-actin level.

Figure S6. Expression of NLS-tagged Cofilin1 did not decrease nuclear F-actin level.

Figure S7. Synthesis of $PEG_{750}$-LANA-JF646 (JF646-LANA).

Figure S8. Mass spectrometry analysis of JF646-LANA.

**Supplementary movies**

Movie 1. F-Actin dynamics visualized using UtrCH-sfGFP in the zebrafish embryo from the 8-cell stage to high stage.

Movie 2–5. F-actin dynamics during chromosome condensation at the 256- (Movie 2), 512- (Movie 3), 1k-cell (Movie 4), and high stage (Movie 5).

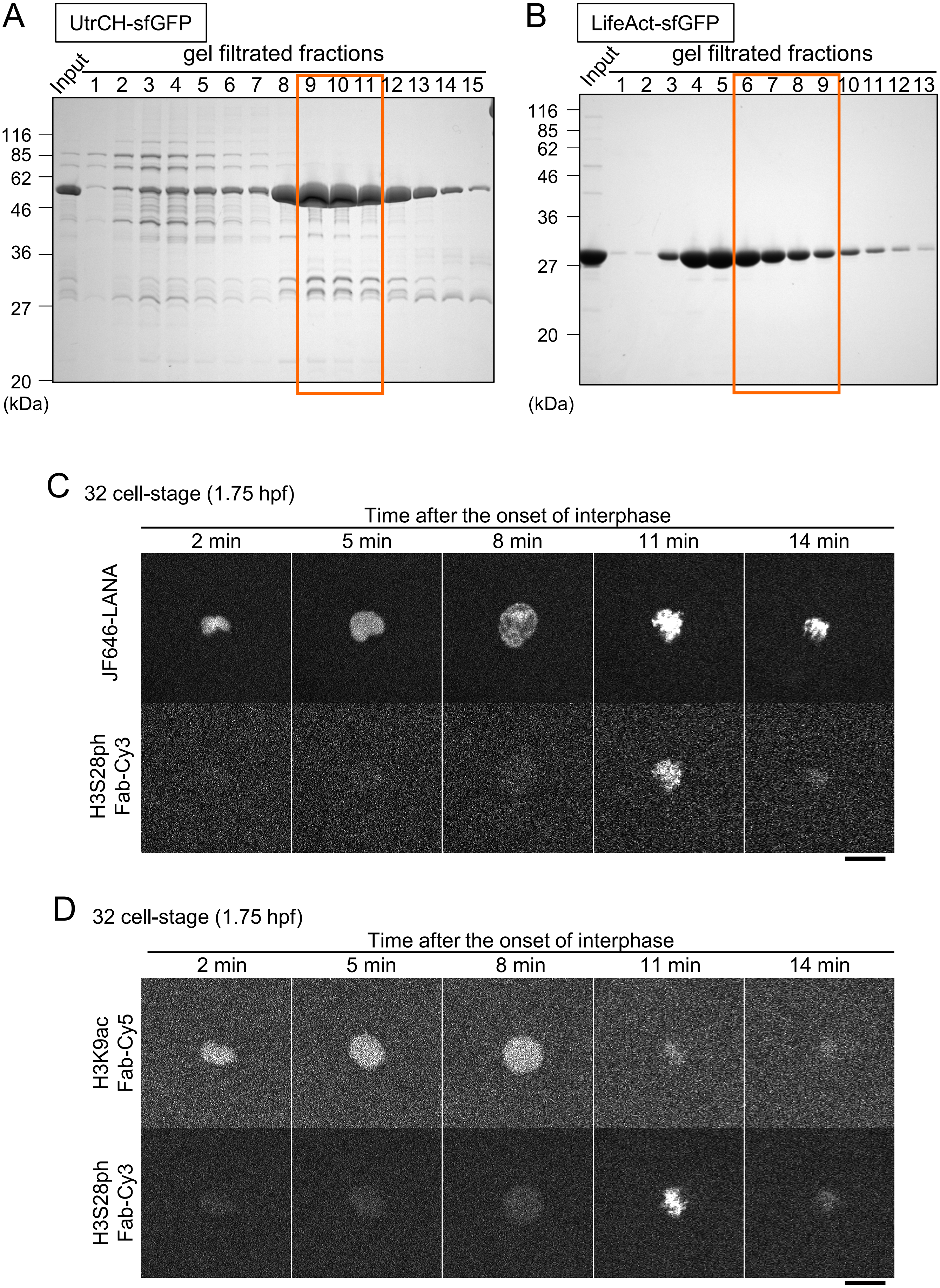
A
UtrCH-sfGFP
gel filtrated fractions
Input
1 2 3 4 5 6 7 8 9 10 11 12 13 14 15
116
85
62
46
36
27
20
(kDa)
B
LifeAct-sfGFP
gel filtrated fractions
Input
1 2 3 4 5 6 7 8 9 10 11 12 13
116
85
62
46
36
27
20
(kDa)
C
32 cell-stage (1.75 hpf)
Time after the onset of interphase
2 min
5 min
8 min
11 min
14 min
JF646-LANA
H3S28ph Fab-Cy3
D
32 cell-stage (1.75 hpf)
Time after the onset of interphase
2 min
5 min
8 min
11 min
14 min
H3K9ac Fab-Cy5
H3S28ph Fab-Cy3



Figure S1

**Figure S1. Actin and chromatin probes.**

(**A** and **B**) Purification of actin probes. UtrCH-sfGFP and LifeAct-sfGFP were expressed in *E. coli* and purified through GST-sepharose followed by GST removal and gel filtration. Fractions though a gel filtration column and inputs were separated on polyacrylamide gels and stained with Coomassie Blue. The positions of size standards are indicated on the left. The fractions #9-11 of UtrCH-sfGFP (**A**) and #6-9 of LifeAct-sfGFP (**B**), as indicated by orange boxes, through gel filtration were pooled and used as actin probes for live imaging. (**C** and **D**) JF646-LANA labels mitotic chromosomes as well as interphase chromatin. Zebrafish embryos were injected with either JF646-LANA (**C**) or H3K9ac Fab-Cy5 (**D**) as a chromatin marker, together with H3S28ph Fab-Cy3 as a mitotic chromosome marker. Confocal sections were acquired every 1 min. Single confocal sections are shown for JF646-LANA (**C**), or H3K9ac Fab-Cy5 (**D**), and H3S28ph Fab-Cy3 at the 32-cell stage. JF646-LANA (**C**) clearly highlights chromatin throughout the cell cycle, with intense signals on condensed chromosomes from prophase (8 min) to prometaphase (11 min) and anaphase (14 min), compared to H3K9ac Fab-Cy5 signals (**D**) that become weaker during mitosis (11 and 14 min). Scale bars: 20 µm

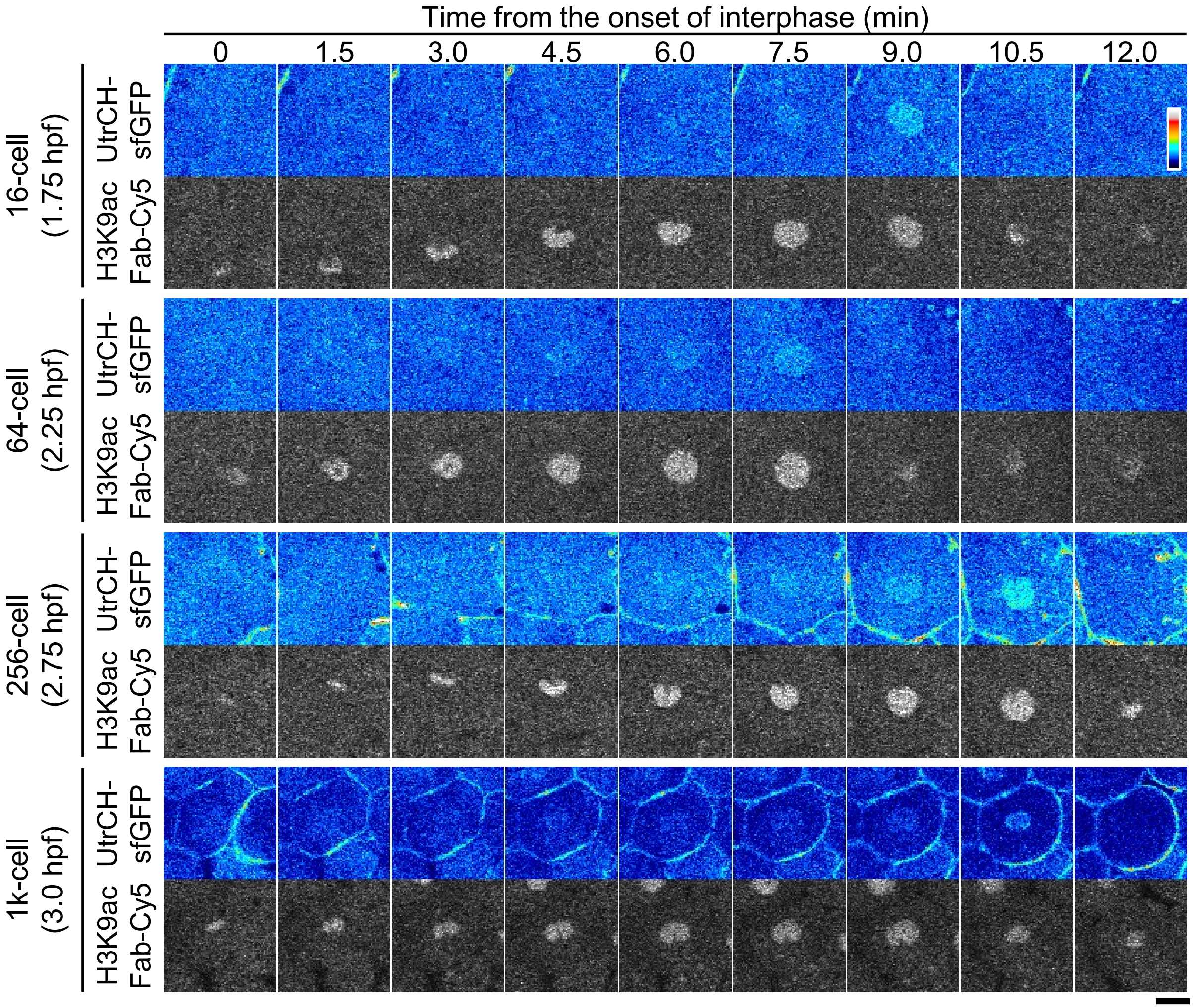


**Figure S2. Actin accumulated in the interphase nuclei at the 16-, 64-, 256-, and 1k-cell stages.**

Zebrafish embryos were injected with UtrCH-sfGFP and H3K9ac Fab-Cy5, and images were acquired using the same procedure as described in Figure 1D and 2A. Single confocal sections of a nucleus at the 16-, 64-, 256-, 1k-cell stages are depicted with the time (min) after the onset of the interphase. Scale bar: 20 μm.



Figure S2

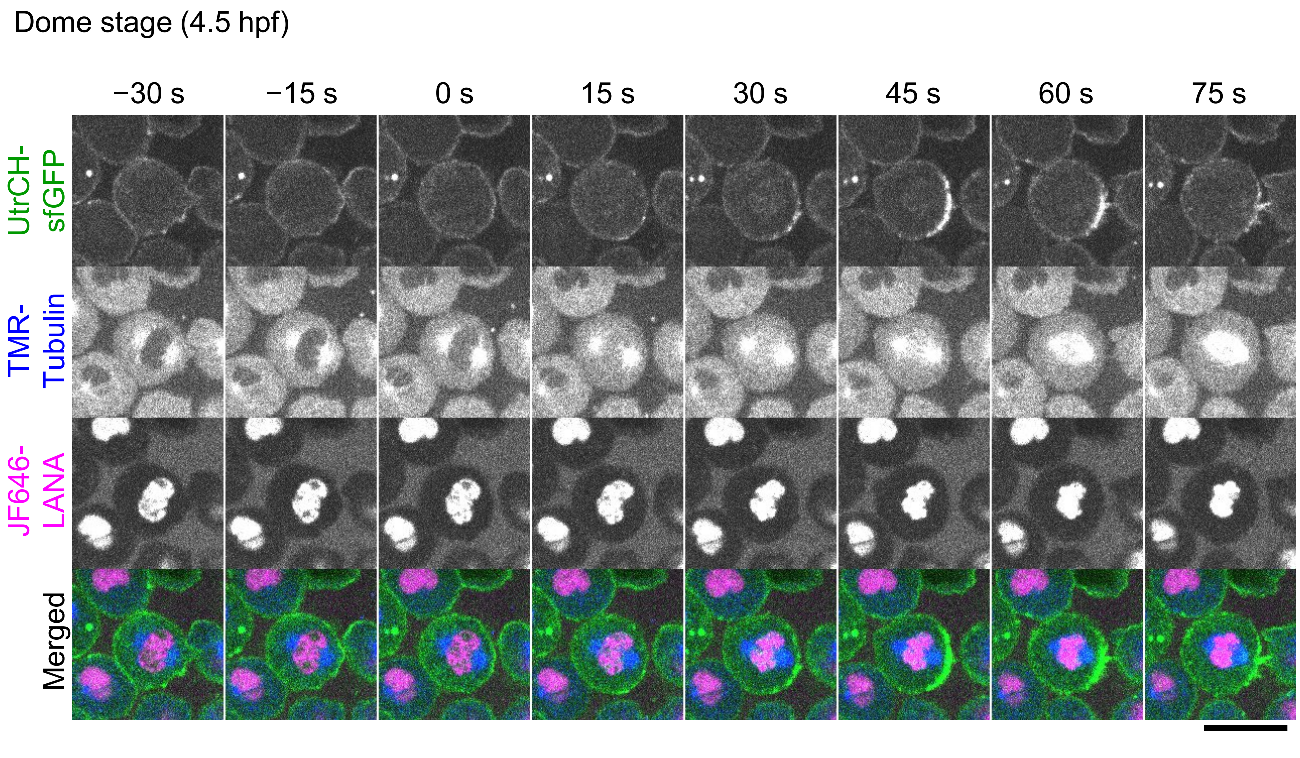


**Figure S3. Weak actin patches in the dome stage embryo.**

Zebrafish embryos were injected with UtrCH-sfGFP, TMR-labeled tubulin, and JF646-LANA. Every 15 s, confocal sections were acquired. Single sections for UtrCH-sfGFP, TMR-tubulin, and JF646-LANA in the dome stage embryo, and their merged images are shown. At this stage, very weak UtrCH-sfGFP signals appeared during prometaphase (from 15 s to 45 s). Scale bar; 20 μm.



Figure S3

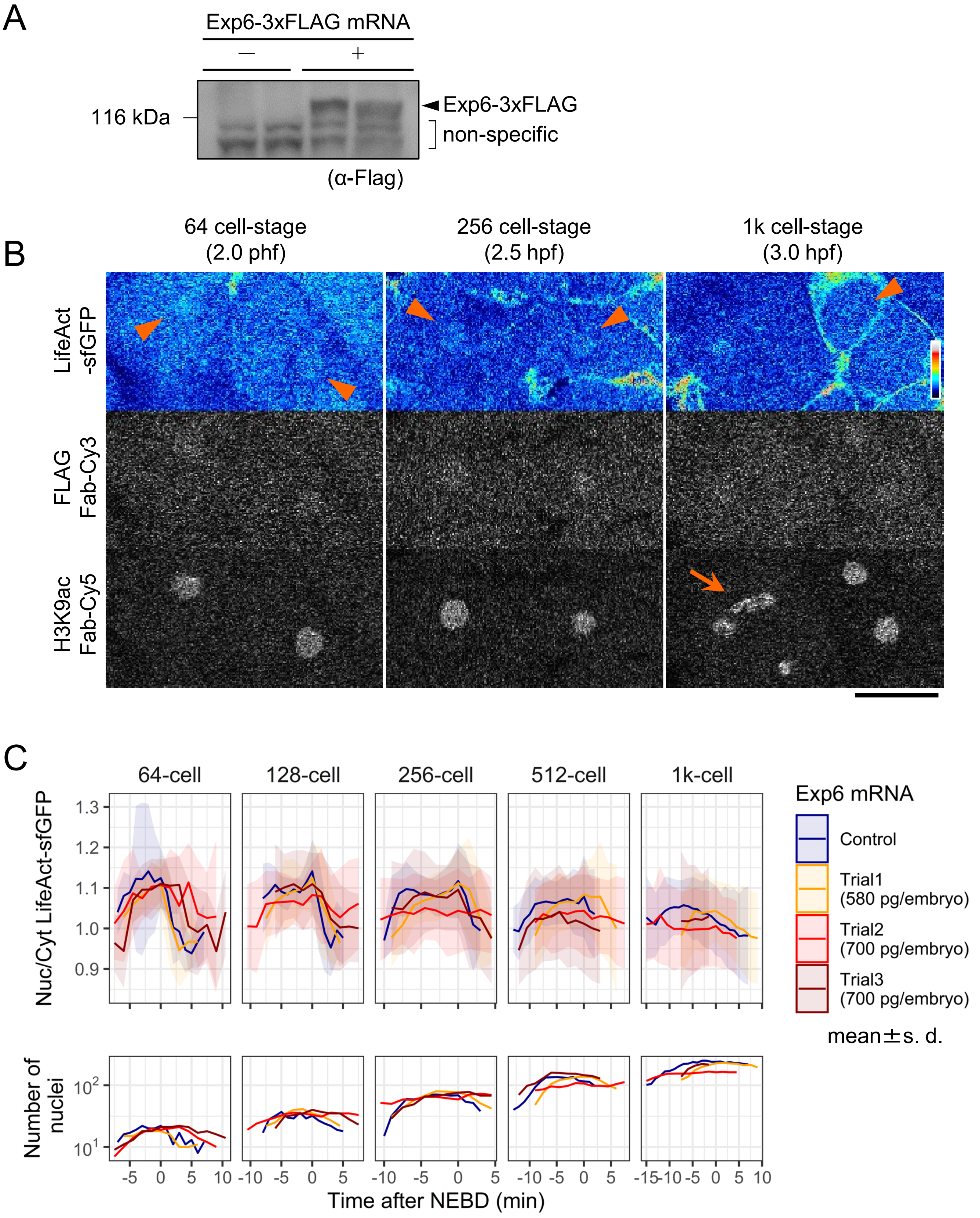
A
Exp6-3xFLAG mRNA
—
+
116 kDa
Exp6-3xFLAG
non-specific
(α-Flag)
B
64 cell-stage
(2.0 phf)
256 cell-stage
(2.5 hpf)
1k cell-stage
(3.0 hpf)
LifeAct
-sfGFP
FLAG
Fab-Cy3
H3K9ac
Fab-Cy5
C
64-cell
128-cell
256-cell
512-cell
1k-cell
Nuc/Cyt LifeAct-sfGFP
Exp6 mRNA
Control
Trial1
(580 pg/embryo)
Trial2
(700 pg/embryo)
Trial3
(700 pg/embryo)
mean±s. d.
Number of
nuclei
Time after NEBD (min)



Figure S4

**Figure S4. Expotin 6 expression did not decrease nuclear actin levels but caused severe defects in cytokinesis.**

(**A**) To express zebrafish exportin 6 (Exp6), embryos were injected with mRNA of 3×FLAG-tagged Exp6, and were allowed to develop for 5.3 h (up to 50% epiboly). After de-yolking, embryos were washed and dissolved in SDS-gel loading buffer. Two lysates from 10 embryos each for control (−) and mRNA-injected (+) were separated on a SDS-polyacrylamide gel for western blotting with anti-FLAG antibody. The positions of the size standard and Exp6-3×FLAG are indicated on the left and right, respectively. (**B**) Zebrafish embryos were injected with mRNA encoding Exp6-3×FLAG, LifeAct-sfGFP, Cy3-labeled anti-FLAG Fab, and Cy5-labeled anti-H3K9ac Fab. Single confocal sections at the 64-, 256-, and 1k-cell stages are shown. LifeAct-sfGFP and FLAG Fab-Cy3 are slightly concentrated in nuclei (arrowheads). Cytokinesis defects were induced and resulted in binucleation (arrow). Scale bar: 50 μm. (**C**) Three independent mRNA injection experiments and one control were performed as in (**B**). The mean values of N/C ratios of LifeAct-sfGFP were plotted with s. d. and the number of nuclei. N/C ratio of LifeAct-sfGFP was not decreased by Exp6-3×FLAG expression.

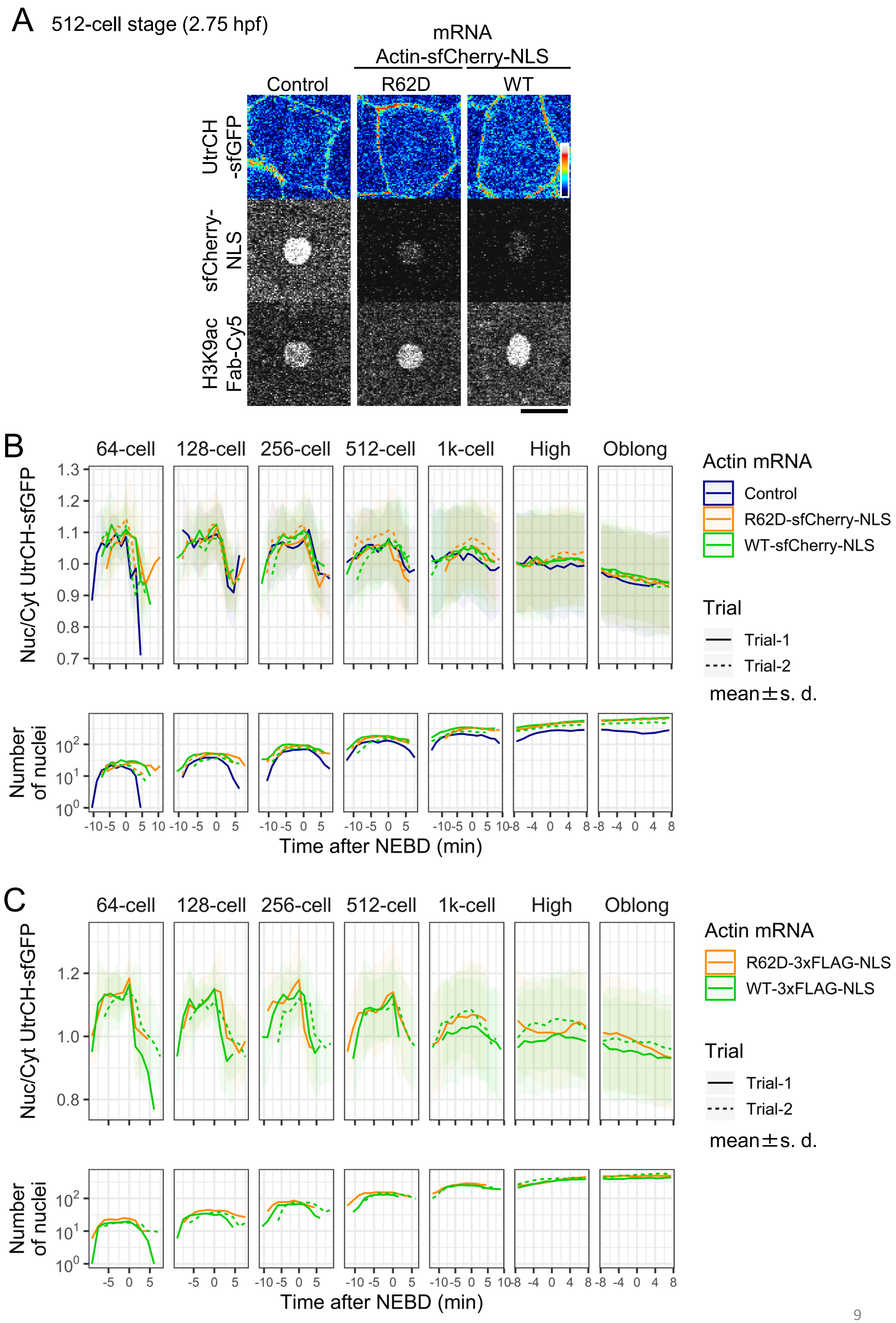
A
512-cell stage (2.75 hpf)
mRNA
Actin-sfCherry-NLS
Control
R62D
WT
UtrCH -sfGFP
sfCherry-NLS
H3K9ac Fab-Cy5
B
64-cell
128-cell
256-cell
512-cell
1k-cell
High
Oblong
Nuc/Cyt UtrCH-sfGFP
Number of nuclei
Time after NEBD (min)
Actin mRNA
Control
R62D-sfCherry-NLS
WT-sfCherry-NLS
Trial
Trial-1
Trial-2
mean±s. d.
C
64-cell
128-cell
256-cell
512-cell
1k-cell
High
Oblong
Nuc/Cyt UtrCH-sfGFP
Number of nuclei
Time after NEBD (min)
Actin mRNA
R62D-3xFLAG-NLS
WT-3xFLAG-NLS
Trial
Trial-1
Trial-2
mean±s. d.



Figure S5

**Figure S5. NLS-tagged polymerization-defective mutant and wild-type actin proteins did not alter nuclear F-actin levels.**

(**A**) Zebrafish embryos were injected with UtrCH-sfGFP and Cy5-H3K9ac Fab followed by mRNA of Actin R62D-sfCherry-NLS or Actin WT-sfCherry-NLS. sfCherry-NLS mRNA was used as a control. Every 90 s, 25 z-stack sections with 4 µm intervals were acquired using a confocal microscope. Single z-sections at the 512-cell stage are shown. Scale bar: 20 µm. (**B**) Two independent experiments Actin R62D-sfCherry-NLS and Actin WT-sfCherry and one control were performed. N/C ratio of UtrCH-sfGFP were measured and the mean values were plotted with s. d. and the number of nuclei. No substantial difference was observed. (**C**) Zebrafish embryos were injected with mRNA of Actin R62D-3×FLAG-NLS or Actin WT-3×FLAG-NLS. UtrCH-sfGFP and Cy5-H3K9ac Fab were co-injected subsequently and 25 z-stack sections with 4 µm intervals were acquired every 90 s using a confocal microscope. N/C ratio of UtrCH-sfGFP were measured and the mean values were plotted with the s. d. and the number of nuclei, as in (**B**). No substantial difference was observed.

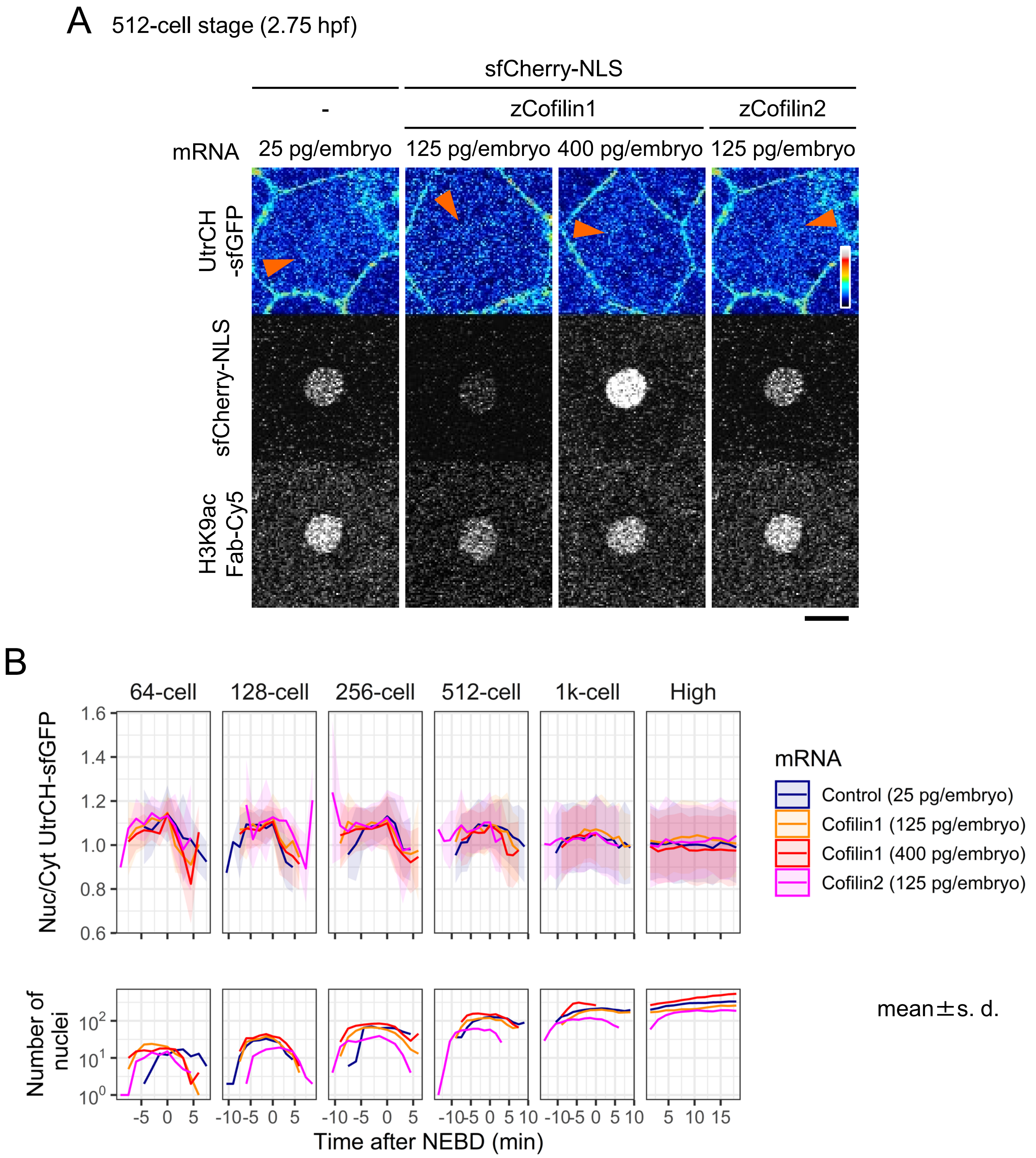
A 512-cell stage (2.75 hpf)
sfCherry-NLS
-
zCofilin1
zCofilin2
mRNA
25 pg/embryo
125 pg/embryo
400 pg/embryo
125 pg/embryo
UtrCH -sfGFP
sfCherry-NLS
H3K9ac Fab-Cy5
B
64-cell
128-cell
256-cell
512-cell
1k-cell
High
Nuc/Cyt UtrCH-sfGFP
Number of nuclei
Time after NEBD (min)
mRNA
Control (25 pg/embryo)
Cofilin1 (125 pg/embryo)
Cofilin1 (400 pg/embryo)
Cofilin2 (125 pg/embryo)
mean±s. d.



Figure S6

**Figure S6. Expression of NLS-tagged cofilin1 did not decrease nuclear F-actin levels.**

Actin depolymerizing factors, zebrafish cofilin1 and cofilin2, each tagged with sfCherry-NLS, were expressed by mRNA injection at 1-cell stage embryos. sfCherry-NLS was used as the control. UtrCH-sfGFP and Cy5-H3K9ac Fab were co-injected and 25-stack confocal images with 4 μm intervals were acquired every 90 s. (**A**) Single confocal sections at the 512-cell stage are shown. Although cofilin1-sfCherry-NLS was accumulated in the nucleus, nuclear UtrCH-sfGFP signals were still detected (arrowheads). Scale bar: 20 μm. (**B**) Mean values of N/C intensity of UtrCH-sfGFP are plotted with the s. d. and the number of nuclei. No substantial difference was observed.

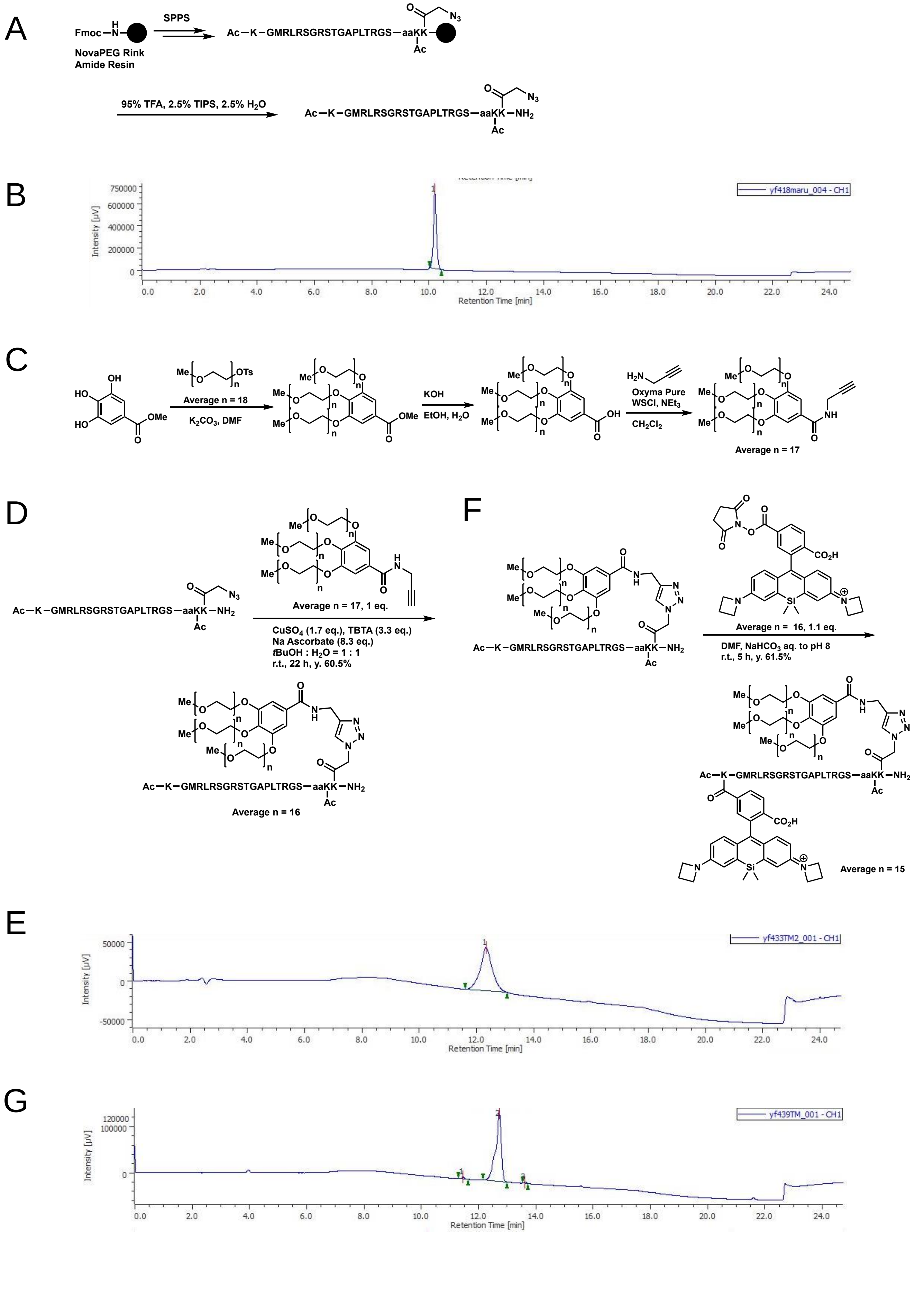

A
Fmoc–N
NovaPEG Rink Amide Resin
SPPS
Ac–K–GMRLRSGRSTGAPLTRGS–aaKK
Ac
N3
95% TFA, 2.5% TIPS, 2.5% H2O
Ac–K–GMRLRSGRSTGAPLTRGS–aaKK–NH2
B
yf418maru_004 - CH1
Intensity [µV]
Retention Time [min]
C
OTs
Average n = 18
K2CO3, DMF
OMe
KOH
EtOH, H2O
OH
Oxyma Pure
WSCl, NEt3
CH2Cl2
Average n = 17
D
Average n = 17, 1 eq.
CuSO4 (1.7 eq.), TBTA (3.3 eq.)
Na Ascorbate (8.3 eq.)
tBuOH : H2O = 1 : 1
r.t., 22 h, y. 60.5%
Average n = 16
F
CO2H
Average n = 16, 1.1 eq.
DMF, NaHCO3 aq. to pH 8
r.t., 5 h, y. 61.5%
Average n = 15
E
yf433TM2_001 - CH1
Intensity [µV]
Retention Time [min]
G
yf439TM_001 - CH1
Intensity [µV]
Retention Time [min]


Figure S7

**Figure S7. Synthesis of $PEG_{750}$-LANA-JF646 (JF646-LANA).**

The reaction schemes for $PEG_{750}$-LANA-JF646 synthesis (**A**, **C**, **D**, and **F**) and analytical HPLC chromatograms (**B**, **E**, and **G**) are shown. (**A**) Synthesis of LANA peptide by sequential addition of Fmoc-amino acid on a NovaPEG-Rink-amide-resin. (**C**) Synthesis of 3,4,5-Tris($PEG_{750}$)-N-(prop-2-yn-1-yl)benzamide. (**D**) Synthesis of $PEG_{750}$-LANA. (**F**) Synthesis of $PEG_{750}$-LANA-JF646. (**B**, **E**, and **G**) Products were analyzed by analytical HPLC using a YMC-Triart Phenyl C18 (4.6 mm I.D. x 150 mm) column with a linear gradient of 2%–90% acetonitrile in 0.1% aqueous TFA over 3–15 min. (**B**) LANA peptide (Retention time: 10.2 min, 22.9 mg, yield 16.9%). (**E**) $PEG_{750}$-LANA (Retention time: 12.3 min, 2.51 mg, 0.436 μmol, yield 60.5%, yellow powder). (**G**) $PEG_{750}$-LANA-JF646 (Retention time: 12.7 min, 1.56 mg, 0.246 μmol, yield 61.5%, powder).

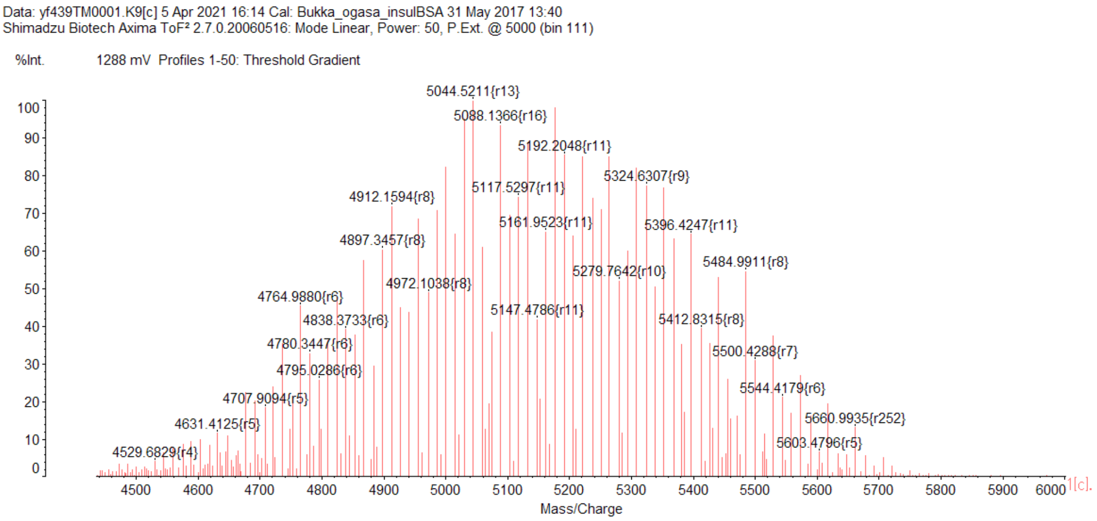


Zoomed view

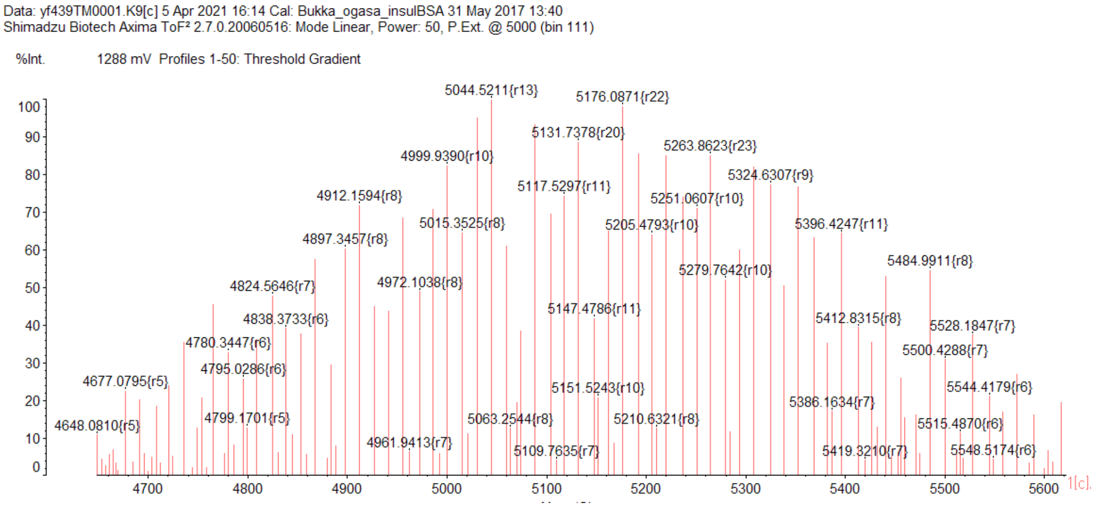


**Figure S8. Mass spectrometry analysis of JF646-LANA.**

Wide (**A**) and zoomed (**B**) views of the MALDI-TOFMS spectrum of $PEG_{750}$-LANA-JF646 are shown. One of the peaks of the highest intensity has *m*/*z* of 5263.9, which corresponds to ethylene glycol n = 15 (Calculated: 5260.9 $[M]^+$ $C_{236}H_{408}N_{45}O_{83}SSi^+$).



Figure S8

**Legends to movies**

**Movie 1. F-actin dynamics visualized using UtrCH-sfGFP in the zebrafish embryo from the 8-cell stage to the high stage.**

Zebrafish embryos were injected with UtrCH-sfGFP and H3K9ac Fab-Cy5. Every 90 s, fluorescence images of the 8-cell stage to the high stage were acquired using a confocal microscope. Single confocal sections of UtrCH-sfGFP signals from the 8-cell stage (1.25 hpf) to high stage (3.5 hpf) are shown.

**Movie 2. F-actin dynamics during chromosome condensation at the 256-cell stage.**

Zebrafish embryos were injected with UtrCH-sfGFP, TMR-labeled tubulin, and JF646-LANA. Every 15 s, fluorescence images of the 256-cell stage (2.5 hpf) were acquired using a confocal microscope. UtrCH-sfGFP patches located in the vicinity of chromosomes from NEBD (0 s) to prometaphase (30 s) and disappeared before metaphase (60s). Time after NEBD is indicated.

**Movie 3. F-actin dynamics during chromosome condensation at the 512-cell stage.**

Zebrafish embryos were injected with UtrCH-sfGFP, TMR-Tubulin, and JF646-LANA. Every 15 s, fluorescence images of the 512-cell stage were acquired using a confocal microscope. UtrCH-sfGFP patches located in the vicinity of chromosomes from prophase (−30 s) to prometaphase (30 s) and disappeared before metaphase (60 s). Time after NEBD is indicated.

**Movie 4. F-actin dynamics during chromosome condensation at the 1k-cell stage.**

Zebrafish embryos were injected with UtrCH-sfGFP, TMR-Tubulin, and JF646-LANA. Every 15 s, fluorescence images of the 1k-cell stage were acquired using a confocal microscope. UtrCH-sfGFP accumulated in the nucleus (−15 s) and remained in the vicinity of chromosomes during prometaphase (0 s to 30 s) and disappeared before metaphase (60 s). Time after NEBD is indicated.

**Movie 5. F-actin dynamics during chromosome condensation at the high stage.** Zebrafish embryos were injected with UtrCH-sfGFP, TMR-Tubulin, and JF646-LANA. Every 15 s, fluorescence images of the high stage (3.3 hpf) were acquired using a confocal microscope. UtrCH-sfGFP patches located in the vicinity of chromosomes at NEBD (0 s) and remained during the prometaphase and disappeared before metaphase (60 s). Time after NEBD is indicated.
